# A Locus Coeruleus- dorsal CA1 dopaminergic circuit modulates memory linking

**DOI:** 10.1101/2021.10.27.466138

**Authors:** Ananya Chowdhury, Alessandro Luchetti, Giselle Fernandes, Daniel Almeida Filho, George Kastellakis, Alexandra Tzilivaki, Erica M Ramirez, Mary Y Tran, Panayiota Poirazi, Alcino J Silva

## Abstract

Individual memories are often linked so that the recall of one triggers the recall of another. For example, contextual memories acquired close in time can be linked, and this is known to depend on temporary increase in excitability that drive the overlap between dorsal CA1 (dCA1) hippocampal ensembles encoding the linked memories. Here, we show that the Locus Coeruleus (LC) cells projecting to dCA1 have a key permissive role in contextual memory linking, without affecting contextual memory formation, and that this effect is mediated by dopamine and not by noradrenaline. Additionally, we found that LC to dCA1 projecting neurons modulate the excitability of dCA1 neurons, and the extent of overlap between dCA1 memory ensembles, as well as the stability of coactivity patterns within these ensembles. This discovery of a neuromodulatory system that specifically affects memory linking without affecting memory formation, reveals a fundamental separation between the brain mechanisms that modulate these two distinct processes.

## Introduction

Most memories are organized into structures, and time is one of the factors underlying the organization of these structures [1–6]. For example, contextual memories encoded close in time (hours, but not days) are linked such that recall of one memory triggers the recall of others acquired within the same temporal window [1, 3]. Furthermore, previous studies showed that the overlap between contextual memory ensembles in dorsal CA1 (dCA1) is greater when contextual memories are linked than when they are not [1]. Any given span of time can include many different events, only some of which may be worth remembering and linking to pre-existing memories. Indeed, there are neuromodulatory systems that use saliency, novelty, and reward to affect the strength of individual memories, and we propose that one or more of these systems may also signal when memory linking should take place. Indiscriminate or inappropriate linking of information may contribute to cognitive deficits associated with neuropsychiatric disorders, including schizophrenia, depression, and post-traumatic stress disorders [7–10].

While CREB activation and subsequent increases in neuronal excitability are thought to trigger the molecular and cellular cascades that lead to memory linking [3, 11–13], it is not known whether there are neuromodulatory mechanisms that specifically regulate memory linking. Neuromodulatory systems have been shown to regulate very specific aspects of behavior, including learning, memory, impulsivity, reward, fear, anxiety, attention, etc. [14–17], and they could also regulate memory linking. Furthermore, the dCA1 has been shown to receive dense projections from several subcortical neuromodulatory nuclei [15, 18], including the serotonergic Raphe Nuclei (RN) [19–22] and the Locus Coeruleus (LC), a major source of both noradrenaline and dopamine [23–28]. Interestingly, although dopaminergic inputs from the VTA project heavily to the ventral CA1 [29], its terminals were shown to be nearly absent in dCA1 [25, 26]. Dopamine has been implicated in the detection of novelty, salience, prediction error, and in the persistence of long-term memory [25, 26, 29–35]. Noradrenaline and serotonin have both been shown to be crucial in detecting the motivational valence and salience of an event [36–39]. Therefore, both the RN and the LC could potentially have a role in modulating memory linking in dCA1.

Here, we demonstrate that dopaminergic fibers from LC to dCA1 are critical for linking contextual memories acquired close in time, but they are not essential for contextual memory formation. The LC modulation of dCA1 affects neuronal excitability in this structure, and is required for the increased overlap between linked memory ensembles in dCA1, as well as for the stability of their coactivity patterns. These results reveal the existence of a neuromodulatory system that specifically regulates the molecular, cellular and circuit mechanism that underlie the formation of temporal memory structures.

## Results

### LC to dorsal CA1 projecting cells are required for contextual memory linking, but not for contextual memory formation

To investigate the involvement of the neuromodulatory areas, LC and RN, in contextual memory linking, we first quantified the percentage of TH+ and 5-HT+ cells activated by a novel context in the LC and RN respectively. Ten min of context exploration triggered a significant increase in cFos+ cells in both the LC (Fig. 1A) and RN (Suppl. Fig 1A) compared to home cage controls. Since these areas responded to contextual exploration, and our contextual memory linking paradigm involves contextual exploration [1], we proceeded to test if the cells projecting to the dorsal hippocampus (dHP) were required for contextual memory linking.

**Figure 1:**
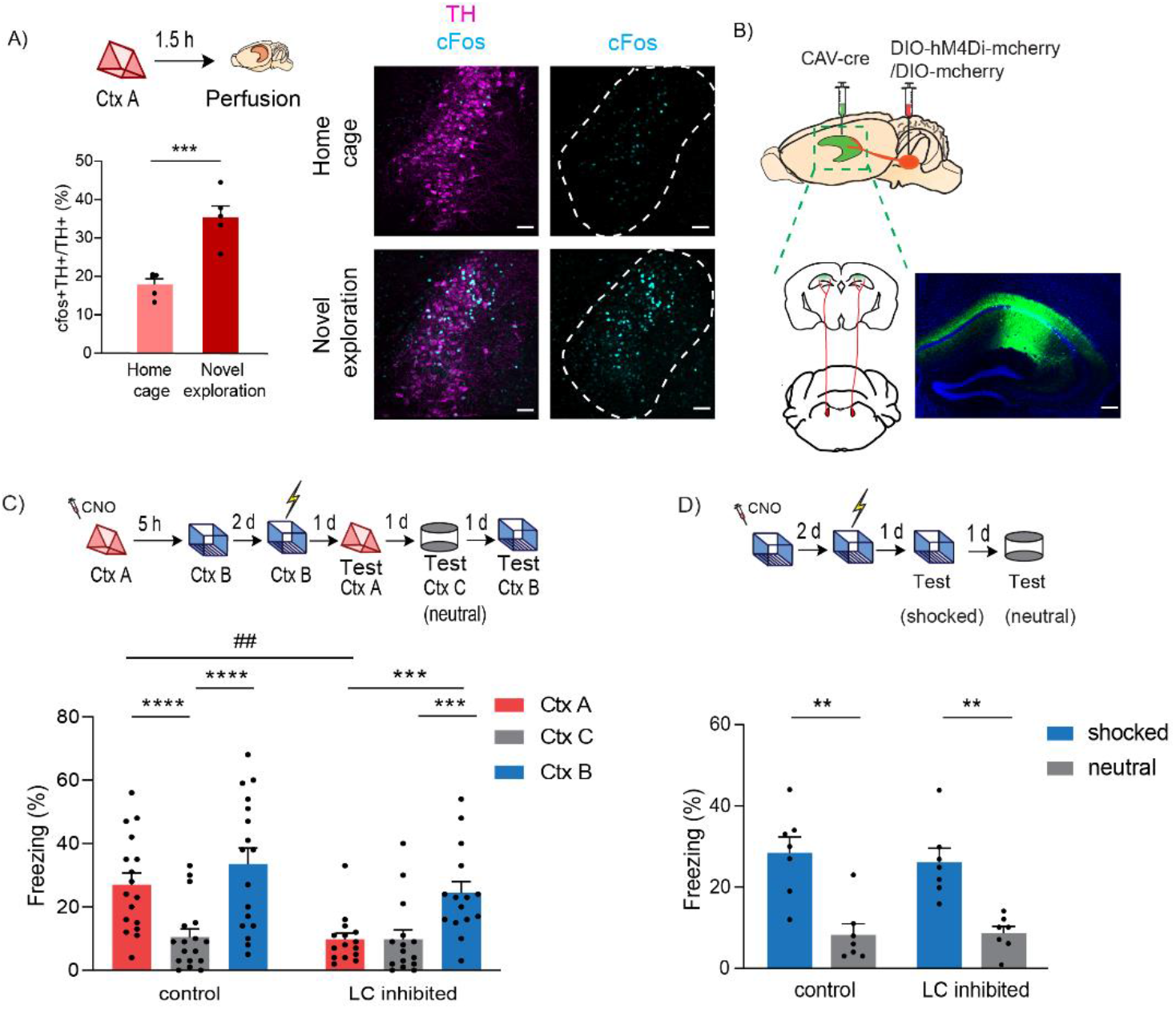
LC to dCA1 projecting cells are required for contextual memory linking, but not for contextual conditioning. A) Exploration of a novel context increased cFos expression in the TH positive cells of LC (unpaired t-test, n=5, ***p<0.001). Example images of TH and cFos staining in LC. Scale bar, 50μm. TH- magenta, cFos- cyan. The LC is outlined. B) Schematics of experimental design for surgery. CAV-cre was injected in dCA1 and DIO-hM4Di-mcherry/DIO-mcherry in LC. Representative image for virus spread in dCA1 estimated with AAV-DIO-GFP injected together with CAV-cre. C) Inhibition of LC cells projecting to dCA1 during exploration of context A impaired contextual memory linking tested with a 5 hours interval. (Control, n=17; LC inhibited, n=15. Two-way repeated measures ANOVA, Sidak post hoc. ##p <0.01, ***p<0.001, ****p<0.0001). Context A- Ctx A, Context B- Ctx B, Context C- Ctx C (neutral). CNO-Clozapine-N-oxide was given to all mice. * is used to depict significance within groups and # is used to show significance between groups for two-way RM ANOVA. D) Inhibition of LC cells projecting to dCA1 did not affect contextual conditioning (Two-way RM ANOVA, Sidak posthoc test; control, n=7; LC inhibited, n=7, **p<0.001). CNO-Clozapine-N-oxide was given to all mice. All results are mean ± s.e.m.

Previous studies have shown that two distinct contextual memories can be linked when they are hours apart, but not when they are a week apart [1]. When two memories are linked, fear paired with one context (context B) is transferred to another neutral context (context A) explored a few hours earlier. During recall, the two linked contexts elicit comparable freezing in mice, while a neutral novel context C triggers significantly lower freezing compared to both the A and B contexts [1].

To investigate the role of LC to dCA1 projecting cells, we used an intersectional viral approach, where a retrograde canine adenovirus (CAV) expressing Cre-recombinase [40, 41] was stereotaxically injected into the dCA1 and AAV-hSyn-DIO-hM4D(Gi)-mCherry (experimental group) or AAV-hSyn-DIO-mCherry (control) [42, 43] was injected into the LC (Fig. 1B). This approach results in the expression of an inhibitory DREADD only in those LC neurons that monosynaptically project to dCA1 (Fig 1B and Suppl. Fig 2A). We used clozapine-N-oxide (CNO, 5mg/kg), administered 30 minutes before a 10-minute exploration of a novel context, to specifically silence the LC to dCA1 projecting neurons (Suppl. Fig 2B and C). Five hours later, the mice were allowed to explore a second novel context (context B), and two days later the mice received a shock immediately upon entering context B. Over the next three days, the mice were tested in context A, context B and in a third novel context (context C - neutral) to test for contextual memory linking, contextual fear conditioning, and contextual generalization, respectively (Fig 1C top). The tests were done on separate days (context B and C recall days were counterbalanced) to minimize interference between tests. The control mice expressing mCherry in LC showed significantly higher freezing in both context A and the shocked context B compared to the neutral context C, (Fig. 1C bottom), demonstrating robust memory linking. Remarkably, the mice expressing DREADD-mCherry in LC froze significantly less in context A compared to context B (shocked context), and freezing in context A was comparable to freezing in the neutral context C. This result demonstrates that inhibiting LC cells projecting to dCA1 during exploration of context A disrupted performance in our contextual memory linking test (Fig. 1C bottom).

Since deficits in contextual memory could confound the interpretation of our contextual memory linking results, we next used the intersectional viral approach just outlined to test whether silencing LC to dCA1 projecting cells affected contextual memory formation (Fig 1D top). The results show that both groups of mice (with and without DREADD-dependent inhibition of the LC to dCA1 projecting neurons) show significantly higher freezing in the shocked context than in the novel context (Fig 1D bottom). This result demonstrates that although inhibiting LC neurons projecting to dCA1 disrupted contextual memory linking, it did not affect the formation of individual contextual memories. Thus, the impairment in contextual memory linking cannot be attributed to deficits in contextual memory formation, a result that revealed a neuromodulatory circuit specific for regulating the linking of memories across time.

### LC to dorsal CA3 projecting cells are required for contextual memory formation

In addition to projecting to dCA1, LC also projects to dCA3 [25, 26, 34]. To investigate the role of LC cells projecting to dCA3 in contextual memory linking and contextual memory formation, we used the same viral approach described above, except that the retrograde CAV expressing Cre-recombinase [40, 41] was stereotaxically injected into dCA3 (Fig. 2A). Then, the mice were CNO treated and tested as described above (Fig. 2B top). The results showed that silencing LC cells projecting to dCA3 disrupted performance in contextual memory linking: the mice showed comparable freezing in contexts A and C and significantly lower freezing than in the shocked context B (Fig. 2B bottom). However, silencing LC cells projecting to dCA3 also disrupted contextual memory formation (Fig. 2C top): The mice with DREADD-dependent inhibition of the LC to dCA3 cells showed a significant reduction of freezing in the shocked context compared to the mCherry controls, and this freezing was indistinguishable from the freezing observed in the neutral context (Fig. 2C bottom). These results were consistent with recent findings showing that optogenetic silencing of LC fibers projecting to dCA3 resulted in a deficit in contextual memory, while silencing of LC projections to dCA1 did not [34].

**Figure 2:**
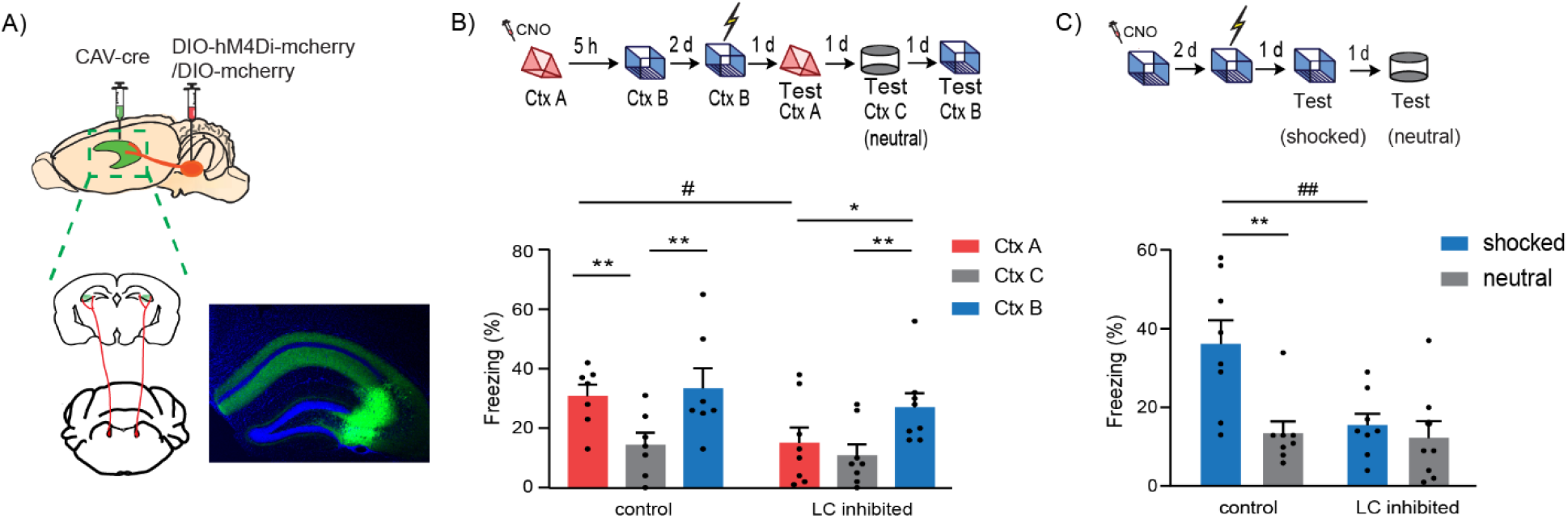
LC to dCA3 projecting cells are required for contextual memory formation. A) Schematics of experimental design for surgery. CAV-cre was injected in dCA3 and DIO- hM4Di-mcherry/DIO-mcherry in LC. Representative image for virus spread in dCA3 estimated with AAV-DIO-GFP injected together with CAV-cre. B) Inhibition of LC cells projecting to dCA3 during exploration of context A impaired performance in contextual memory linking tested with a 5 hours interval. (Control, n=7; LC inhibited, n=8. Two-way repeated measures ANOVA, Sidak post hoc. #p <0.05, *p < 0.05, **p<0.01). * is used to depict significance within groups and # is used to show significance between groups for two-way RM ANOVA. C) Inhibition of LC cells projecting to dCA3 impaired contextual conditioning (control, n=8; LC inhibited, n=8; Two-way RM ANOVA, Sidak posthoc test; **p<0.001, ##p<0.01). * is used to depict significance within groups and # is used to show significance between groups for two-way RM ANOVA. All results are mean ± s.e.m.

### RN to dorsal CA1 projecting cells are not required for contextual memory linking

Next, we tested whether RN cells projecting to dCA1 are necessary for contextual memory linking using a similar approach to the one outlined above: the retrograde CAV virus expressing Cre-recombinase was stereotaxically injected into the dCA1, and AAV-hSyn-DIO-hM4D(Gi)-mCherry (experimental group) or AAV-hSyn-DIO-mCherry (controls) were injected into the RN (Suppl. Fig 3A). As described above, the mice in both groups received a CNO injection 30 min before exploration of context A, and five hours later they were allowed to explore context B. Two days after that, the mice received a shock immediately upon entering context B. As before, over the next three days, the mice were tested in context A, context B and in a third novel context (context C; Suppl. Fig 3B top). Both groups of mice showed significantly higher freezing in context A and B than in the neutral context C, (Suppl. Fig 3B bottom), demonstrating robust contextual memory linking. This result shows that inhibiting RN cells projecting to dCA1 does not impair contextual memory linking.

All together the results presented above indicate that while LC to dCA1 projecting neurons are specifically involved in contextual memory linking, LC to dCA3 projecting neurons are required for contextual memory formation, and RN cells projecting to dCA1 are not required for either of these two processes. These findings revealed that memory linking and memory formation can be independently regulated, and that LC to dCA1 projecting cells have a critical and specific role in this process.

### LC modulates neuronal excitability in dCA1

Memory formation is known to induce transient increases in neuronal excitability [13, 44–49], biasing the allocation of subsequent memories to the same neuronal ensembles [1, 3–5, 50–52] and thus linking the two memories [1, 3, 52]. Hence, we reasoned that the LC may control contextual memory linking by modulating the persistence of dCA1 neuronal excitability.

We tested this hypothesis using the same intersectional viral strategy described above, and systemically administered CNO (5mg/kg) 30 minutes before the mice explored a novel context (context A). The excitability of dCA1 neurons was measured 5 hours after exposure to context A (Fig. 3A). We found that inhibiting LC to dCA1 projecting cells reduced the firing rate of dCA1 neurons in response to increasing steps of current injection, indicating a reduction in neuronal excitability (Fig. 3B and C). This showed that LC to dCA1 projecting cells modulate dCA1 neuronal excitability triggered by novel context exploration, a cellular mechanism previously proposed to be critical for contextual memory linking [1]. In contrast, there were no changes in the resting membrane potential (RMP) and input resistance (R_in_) of dCA1 neurons (Suppl. Table 1). These results showed that the inhibition of LC to dCA1 projecting cells modulates the learning-induced dCA1 neuronal intrinsic excitability, which is consistent with the impact of this manipulation on contextual memory linking.

**Figure 3:**
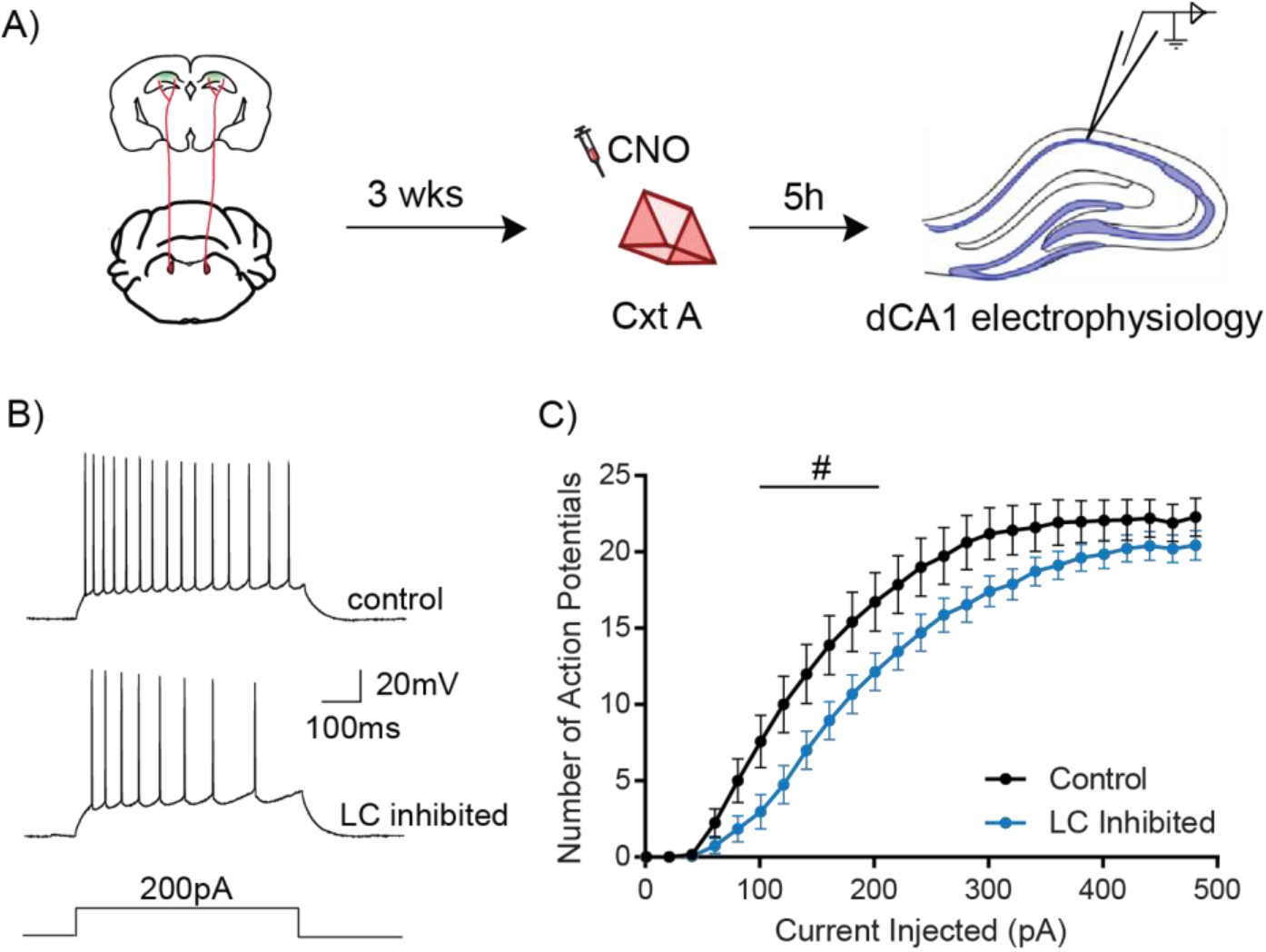
LC to dCA1 inhibition decreases novelty induced increases in neuronal excitability. A) Experimental design for measuring dCA1 neuronal excitability 5 hours after context exploration with chemogenetic inhibition of dCA1 projecting LC neurons. B) Representative traces showing adaptive firing responses to a 200pA current injection in dCA1 pyramidal neurons. C) Inhibition of LC cells projecting to dCA1 during context exploration reduced the firing rate of dCA1 neurons 5 hours later (Two-way repeated measures ANOVA; control n=15, LC inhibited n=15, #p<0.05). # is used to show significance between groups for two-way RM ANOVA. All results are mean + s.e.m.

### LC modulates the overlap of contextual memory ensembles in dCA1

Studies of memory linking showed that the overlap between memory ensembles was critical for this process [1, 3, 52]. Specifically, previous results indicated that contextual memory linking depends on the overlap between dCA1 contextual memory ensembles, since ensembles of linked memories showed more overlap than unlinked ones, and two unlinked memories could be linked simply by artificially increasing the overlap between their dCA1 memory ensembles [1]. These results, together with excitability findings presented above, suggest that LC to dCA1 projecting cells may modulate contextual memory linking by regulating the overlap between contextual memory ensembles in the dCA1.

To test this hypothesis, we injected a GCaMP6f AAV virus (AAV.Syn.GCaMP6f.WPRE.SV40) into dCA1 and recorded neuronal calcium ensemble activity in dCA1 with head mounted fluorescent microscopes (miniscopes) [1, 53–55] while mice explored two different novel contexts (A and B) separated by 5 hours (Fig 4A). Using the intersectional viral strategy described above, we inhibited LC neurons projecting to dCA1 while the mice explored the first context (context A), and then measured the overlap between the neuronal populations recorded during the two contextual exposures (Fig. 4A). Compared to controls, the mice with inhibition of LC neurons projecting to dCA1 showed a significant reduction in the overlap between contextual memory ensembles activated during exploration of contexts A and B (Fig 4B). Importantly, there was no significant difference between the active population of cells detected with and without inhibition of LC neurons projecting to dCA1 (Suppl. Fig 4A). Studies of c-Fos expression also confirmed that inhibition of LC neurons projecting to dCA1 did not affect the overall activation of dCA1 neurons during contextual exploration (Suppl. Fig 4B).

**Figure 4:**
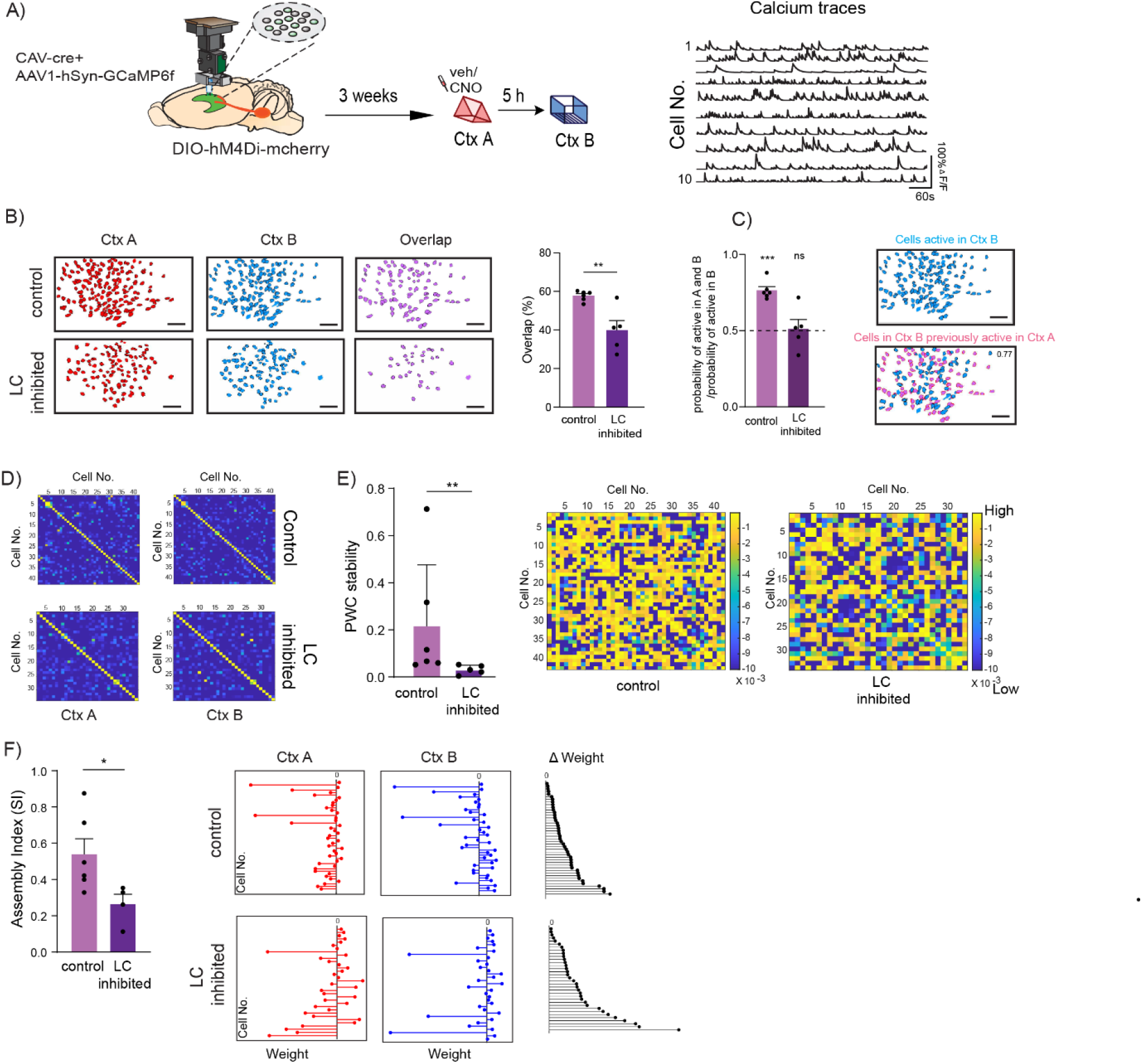
LC to dCA1 inhibition decreases the overlap between dCA1 memory ensembles and affects their firing properties. A) Schematics for miniscope setup and calcium signal imaging in dCA1. CAV-cre was injected in dCA1 and DIO-hM4Di-mcherry was injected in LC. Veh or CNO was injected in the control and LC-inhibited group, respectively. Example calcium traces. B) Inhibition of LC neurons projecting to dCA1 reduced the percentage overlap between memory ensembles encoding contexts explored 5 hours apart (Control n=6, and LC inhibited n=5; unpaired t-test, **p<0.01). Percentage overlap was calculated as neurons active in both A and B over total cells active in A and B. Representative plots of active neurons in contexts A and B and neuronal overlap between different conditions. Scale bars, 50 μm. C) Inhibition of LC neurons projecting to dCA1 reduced the likelihood to chance levels that a cell active in context B had previously been active in context A (Control n=6, and LC inhibited n=5; One-sample t-test over 0.5 as chance level, ***p< 0.001). D) Example maps of pairwise temporal correlations between the activity of all overlapping neurons in context A and B. E) Within the overlapping neuronal population, inhibition of the LC neurons projecting to dCA1 reduced the stability of the coactivity maps between the two contexts visited (Control n=6, and LC inhibited n=5; Mann Whitney test, **p<0.01). Example PWC stability maps. F) Inhibition of the LC neurons projecting to dCA1 reduced the stability of the dCA1 assemblies detected within the overlapping neuronal population (Control n=6, and LC inhibited n=4; unpaired t-test, *p<0.05). Representative images for weight distribution of the assemblies detected for context A (red), context B (blue), or delta between the two weights (black) with all neurons sorted in the same order in all 3 graphs. All results are mean + s.e.m.

Next, we tested whether inhibition of LC cells projecting to dCA1 affected the probability of finding dCA1 cells active in both contexts A and B. The results showed that this probability is above chance levels in control mice, but not in mice with inhibition of LC cells projecting to dCA1(Fig. 4C), a result consistent with our dCA1 memory ensemble overlap findings.

Since inhibition of LC cells projecting to dCA1 affected the probability of finding dCA1 cells active in both contexts A and B, we next studied the impact of this inhibition on co-activity patterns between dCA1 neurons [56, 57]. Groups of neurons with synchronized activity have been suggested to encode task-relevant information, in a number of brain structures, including the hippocampus, cortical, and subcortical regions [56–62]. However, the significance of such co-activity patterns during memory linking in dCA1 is unknown. We first defined coactivity maps for each context, as maps of the pairwise temporal correlations between the activity of all dCA1 neurons that were active in both contexts (Fig. 4D). We found that, compared to controls, inhibition of LC cells projecting to dCA1 significantly decreases the overall stability of the coactivity maps between the two contexts visited (Fig. 4E). To further narrow down our analysis, we identified cell assemblies (subsets of dCA1 neurons that significantly fire together) within the neurons that were active in both contexts [63]. Consistently, inhibition of the LC cells projecting to dCA1 also decreased the stability of these dCA1 cell assemblies (Fig. 4F). This indicated that inhibition of LC cells projecting to dCA1 disrupted the partnerships between dCA1 neurons firing together during the exploration of context A compared to exploration of context B. We found that mean firing rates of the neurons that were activated in both contexts were not significantly affected by inhibition of the LC cells projecting to dCA1 (Suppl. Fig. 4 C and D). This inhibition also did not affect the total number of assemblies detected (Suppl. Fig. E and F), or the mean pair-wise correlations (PWC) (Suppl. Fig. G and H) in either session.

Together, these results strongly support the hypothesis that LC modulates contextual memory linking by biasing the co-allocation of contextual memories acquired close in time to overlapping neuronal ensembles in dCA1. Additionally, our findings suggested that the coactivity patterns between dCA1 neurons may also play a role in memory linking, since LC cells projecting to dCA1 were also shown to regulate the stability of assembly dynamics.

### Modeling the impact of LC modulation of dCA1 neuronal ensembles

Our *in vivo* experiments showed that the LC cells projecting to dCA1 were critical for the increase of neuronal excitability observed after learning. This persistent increase in excitability is thought to underlie the increased overlap between the ensembles of memories separated by hours, thereby linking those memories [1, 5].

To assess whether the LC-mediated increase in excitability is sufficient to account for the neuronal population overlap observed in our experiments, we used a computational modeling approach. Specifically, we adapted a previously published network model [64, 65], which consists of simplified spiking neurons (soma plus a few dendritic compartments) for both excitatory and inhibitory (soma-targeting and dendrite-targeting interneurons) cell types as well as calcium-dependent and protein-dependent Hebbian plasticity (Fig. 5A). In line with the experiments, we simulated the encoding of two distinct contextual memories (A and B) via the activation of two input populations that project to the excitatory model neurons (Fig 5A). Next, we simulated the LC-induced increase in excitability seen in our experiments (Fig. 3B and C) as modulation of the adaptation (AHP) current (Fig. 5B). A delay period of 5 hours was introduced between the encoding of the two memories, during which plasticity processes take place. We simulated two conditions, control and LC-dCA1 inhibited, and analyzed the properties of the entire neuronal population after encoding the two memories. The excitability of the neurons encoding context A was increased under the control condition, but not when LC neurons projecting to dCA1 were inhibited. We found that inhibition of LC input to the network model causes a large drop in the overlap between the populations that are active (i.e. have a firing frequency>10Hz) [64] during the encoding of both context A and context B (Fig. 5C), in agreement with our experimental observations. This large drop in the population overlap is not accompanied by significant changes in the sizes of the populations that are activated by either memory (Fig. 5D). These simulation results agree with our experimental observations, where inhibition of LC input to dCA1 does not affect the overall activation of dCA1 neurons during exploration while disrupting the overlap between two memories close in time.

**Figure 5:**
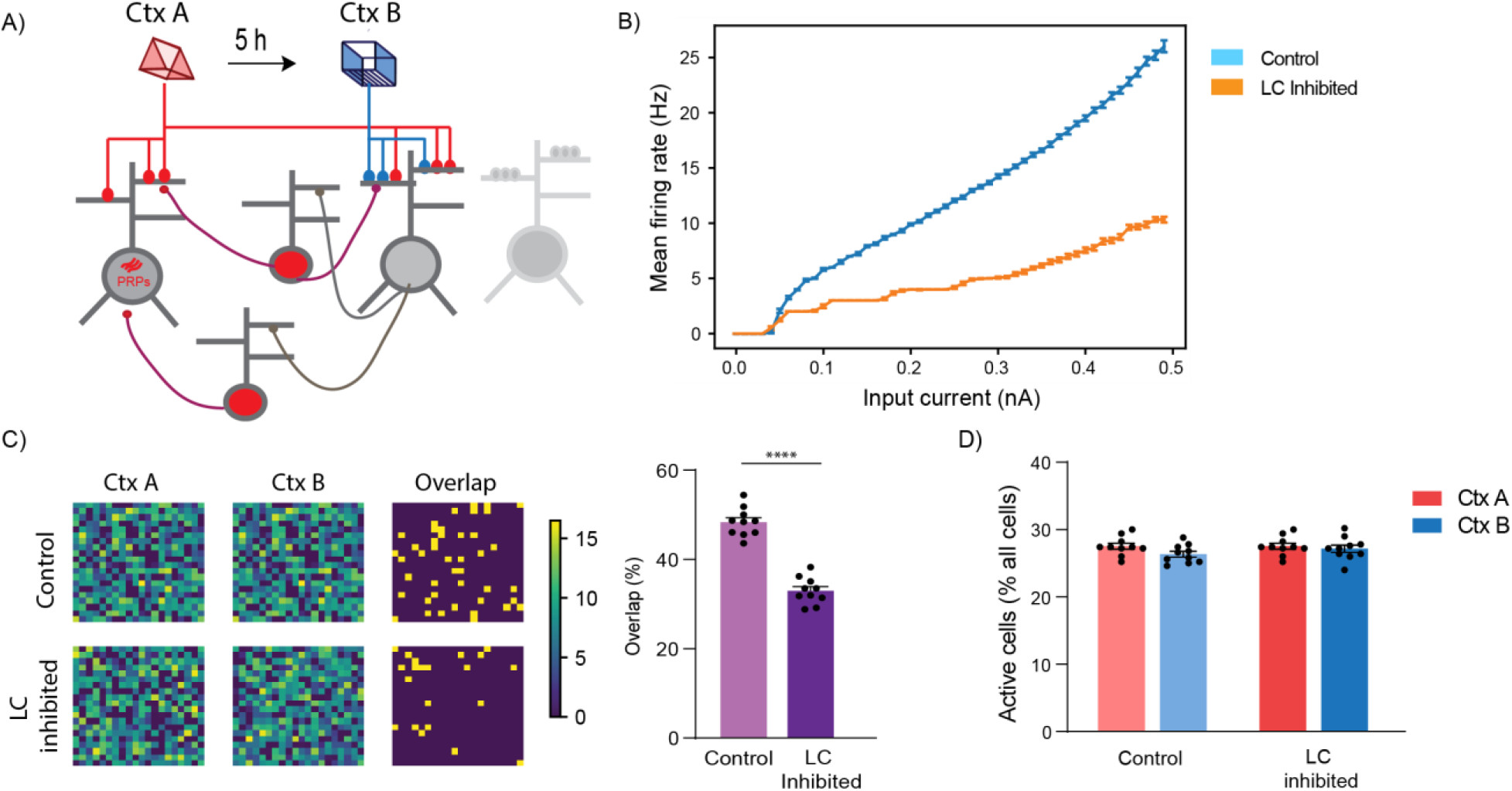
Modeling LC to dCA1 inhibition using a spiking network model. A) Conceptual diagram of the spiking network model. The model includes excitatory neurons (gray) and subpopulations of interneurons (red). Synaptic inputs representing different memories terminate in overlapping dendrites. Two novel contexts are simulated as memories A and B, separated by 5h. B) Firing rate of neurons when current input is applied directly to the somatic compartment of the 2-stage neurons under control (blue) and LC inhibition (orange) condition. Under conditions of LC blocking, the excitability of the neurons does not increase, n=50. C) Simulation of LC to dCA1 inhibition resulted in a reduction of the overlapping neuronal population. Percentage overlap was calculated as neurons activated during both context A and B over the total active neurons in A and B. n=10 simulation trials, unpaired t-test, ****p<0.0001. The contour plots show population activation during encoding of memories in context A and B. The third column indicates the neurons which were active (ff > 10Hz) during both memories. D) The sizes of activated populations (number of neurons with ff>10Hz) during the encoding of Ctx A and Ctx B, under different conditions. All results are mean + s.e.m.

Overall, our modeling simulations confirmed that the modulation of excitability during the encoding of two separate memories combined with Hebbian plasticity rules can shape the overlap of populations activated by both memories without affecting their size, which is in agreement with our experimental hypotheses. They also indicate that mechanisms which affect the adaptation currents could be mediating the effect of the LC neuromodulatory input [66–68].

### Dopamine D1/D5 receptors in dCA1 modulate contextual memory linking

The results presented above demonstrate that inhibition of LC cells projecting to dCA1 disrupt both memory ensemble dCA1 overlap and contextual memory linking. Previous studies showed that LC cells projecting to dCA1 co-release both noradrenaline and dopamine, and that these neuromodulators have differential roles in memory processes [25–28]. To determine which of those two neuromodulators mediate the effects on contextual memory linking, we implanted cannulas bilaterally into the dCA1 of mice, and treated them with either a dopamine D1/D5 receptor antagonist (SCH23390) [25, 26, 69] or with a noradrenaline β-adrenergic receptor antagonist (propranolol) [25, 26] 20 minutes before the mice explored context A in our contextual memory linking test. We found that the dopamine D1/D5 receptor antagonist disrupted contextual memory linking in a dose dependent manner (Fig. 6A), while the noradrenaline β-adrenergic receptor antagonist did not (Fig. 6C). Importantly, in another set of experiments, we confirmed that the doses of the dopamine D1/D5 antagonist that disrupted contextual memory linking did not affect contextual memory itself (Fig. 6B), demonstrating that the effects of the D1/D5 antagonist are specific to memory linking. Together, these results indicate that dopamine (not noradrenaline) signaling in the dCA1 is critical for contextual memory linking.

**Figure 6:**
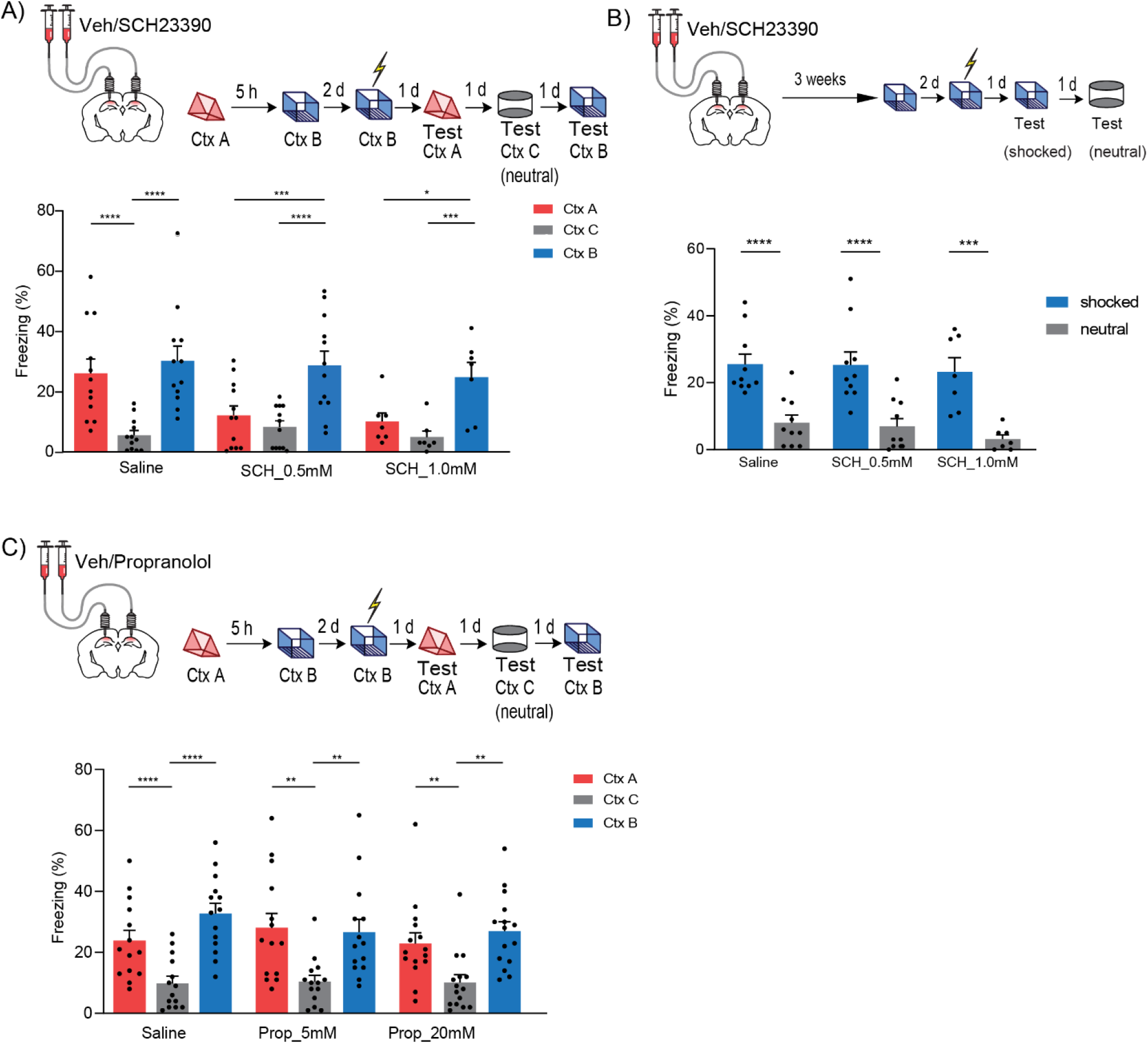
Dopamine D1/D5 receptors in dCA1 modulate contextual memory linking. A) Inhibition of Dopamine D1/D5 receptors in dCA1 during context A disrupted contextual memory linking. (Saline, n=12; SCH 0.5 mM, n=12; SCH 1.0 mM, n=7; two-way repeated measures ANOVA, Sidak post hoc, *p<0.05, **p<0.01, ***p<0.001, ****p<0.0001) B) Inhibition of Dopamine D1/D5 receptors in dCA1 did not affect contextual memory formation at the doses used. (Saline, n=10; SCH 0.5 mM, n=10; SCH 1.0 mM, n=7; two-way repeated measures ANOVA, Sidak post hoc, ***p<0.001, ****p<0.0001). C) Inhibition of β-adrenergic receptors in dCA1 did not affect contextual memory linking at the doses used. (Saline, n=14; Prop 5 mM, n=14; Prop 20 mM, n=15; two-way repeated measures ANOVA, Sidak post hoc, **p<0.01, ****p<0.0001) All results are mean + s.e.m.

### Optogenetic D1 receptor activation in dCA1 rescues linking deficits caused by inhibition of LC cells projecting to dCA1

The results presented above suggested that dopaminergic modulation of the dCA1 by the LC is critical for contextual memory linking. Therefore, these results predict that the deficit in contextual memory linking, caused by inhibition of LC cells projecting to dCA1, could be rescued by activating D1 receptors in dCA1 neurons during contextual memory linking (Fig. 7A). To test this hypothesis, we used a multi-intersectional viral strategy, where we injected retrograde CAV virus expressing Flp-recombinase in dCA1 [70] and Flp-dependent (Frt) AAV-fDIO-hM4i-DREADD-mCherry [71] in the LC to chemogenetically inhibit the LC cells projecting to dCA1. In addition, to activate D1 receptor signaling in dCA1, we injected a cocktail of AAV-hSyn-cre and AAV-DIO-optoD1-GFP [72, 73] in dCA1 while implanting an optic cannula (Fig. 7B). This cre-DIO system, under the transcriptional control of the pan-neuronal synapsin promoter, directs expression to both excitatory and inhibitory neurons [74–76]. Together, this allowed us to chemogenetically inhibit the LC neurons projecting to dCA1 with the Flp-fDIO system, while optogenetically activating D1 receptor signaling in a subset of dCA1 neurons (Fig. 7A and B). We confirmed that the Flp-fDIO and cre-DIO were mutually exclusive: one genetic system did not affect the other [71] (Suppl. Fig 6A and B).

**Figure 7:**
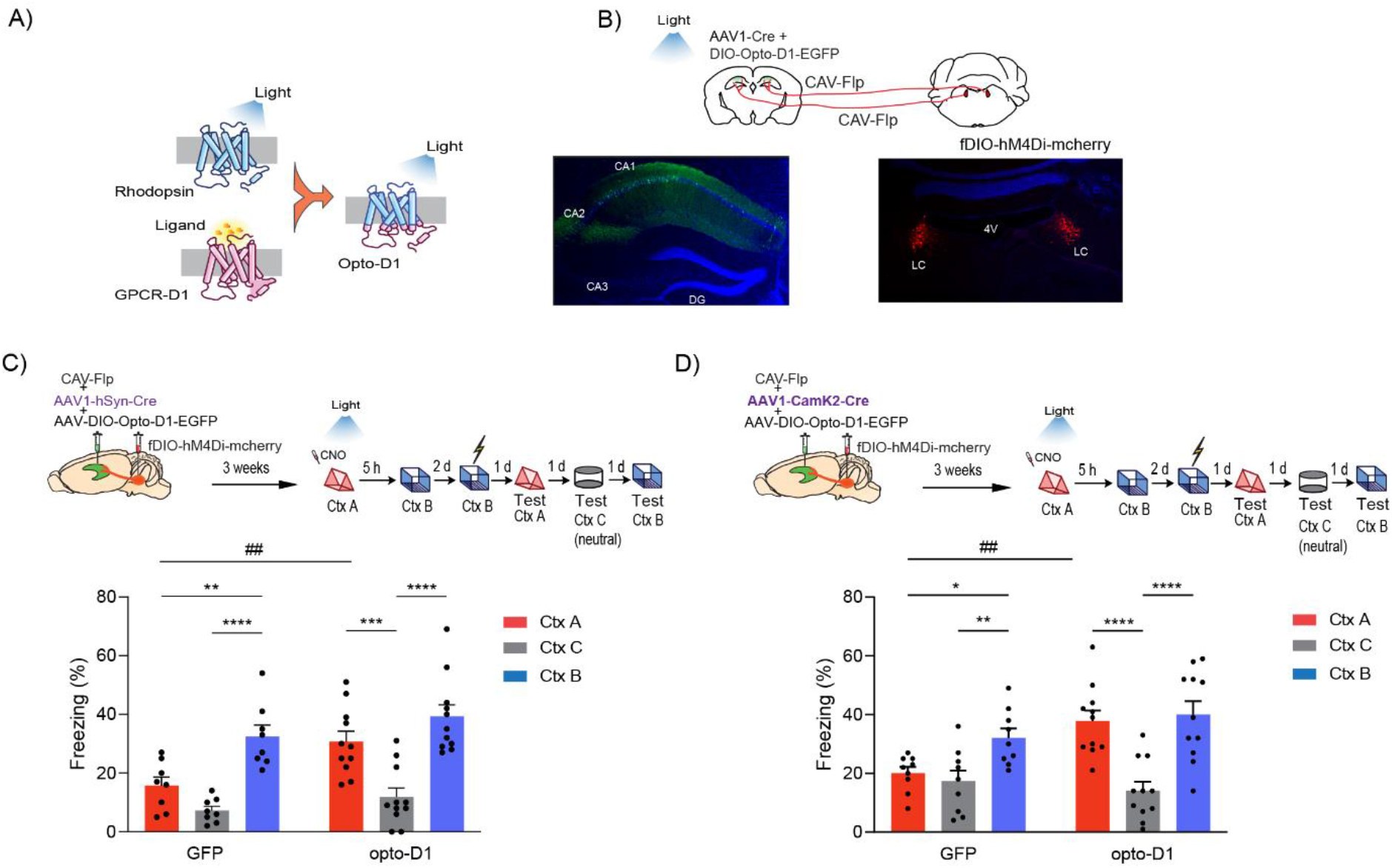
Optogenetic D1 receptor activation in dCA1 rescues linking deficits caused by inhibition of LC-dCA1. A) Schematics of the Opto-D1 construct. B) Schematics of experimental design. AAV-cre with DIO-Opto-D1 was injected in dCA1 to activate D1 signaling, while CAV-FLP was injected in the same location of dCA1 to be retrogradely taken up by cells projecting to this region. Flippase-dependent fDIO-hM4Di-mcherry was injected into LC to silence the neurons projecting to dCA1. C) Optogenetic activation of D1 receptor signaling in a fraction of all cell types in dCA1 rescued the contextual memory linking deficit caused by chemogenetic inhibition of LC cells projecting to dCA1. (GFP, n=8; opto-D1, n=11; two-way repeated measures ANOVA, Sidak post hoc,*p<0.05, **p<0.01, ***p<0.001, ****p<0.0001, ##p<0.01). * is used to depict significance within groups and # is used to show significance between groups for two-way RM ANOVA. D) Optogenetic activation of D1 receptor signaling in a fraction of excitatory cells in dCA1 rescued the contextual memory linking deficit caused by chemogenetic inhibition of LC cells projecting to dCA1. (GFP, n=9; opto-D1, n=11; two-way repeated measures ANOVA, Sidak post hoc,*p<0.05, **p<0.01, ****p<0.0001, ##p<0.01). * is used to depict significance within groups and # is used to show significance between groups for two-way RM ANOVA. All results are mean + s.e.m.

We chemogenetically inhibited LC cells projecting to dCA1 with CNO administered 30 minutes before exploration of the first context A, and optogenetically activated D1 receptors with blue light (473nm, 50s on-10s off, 10mW) while the mice explored context A for 10 minutes, after which they were returned to their home cages. Five hours later, the mice were allowed to explore the second context (context B) without CNO infusion or blue light illumination. Two days later, the mice were returned to context B, where they received an immediate shock. Over the next three days, they were tested in context A, context B and in a third neutral context (context C) (Fig. 7C top). We confirmed that the control mice (with the AAV-DIO-GFP instead of opto-D1, in dCA1) showed a deficit in contextual memory linking, displaying freezing levels in context A that were significantly lower than in the shocked context B, and comparable to the neutral context C (Fig 7C bottom). These results confirm that LC to dCA1 inhibition carried out with CAV-Flp:fDIO-Gi impaired contextual memory linking just as in the experiments described above with the CAV-cre:DIO-Gi system. In contrast, the mice with opto-D1 activation showed significantly higher freezing in context A compared to freezing in the neutral context C, and freezing in context A was similar to freezing in the shocked context B, demonstrating normal contextual memory linking (Fig. 7C bottom). Altogether, these results show that optogenetic activation of dCA1 D1 receptors, specifically during exploration of context A, is sufficient to rescue the loss of memory linking caused by inhibition of LC cells projecting to dCA1.

We next repeated the above experiment while restricting the expression of opto-D1 to excitatory neurons by using the alphaCaMKII promoter [74, 77]. We replaced the AAV-hSyn-cre with the AAV-CaMKII-cre virus (Fig. 7D top), so that the recombinase expression would be restricted to excitatory neurons. The results confirmed that, as before, control mice expressing GFP, instead of opto-D1, showed a deficit in contextual memory linking as expected when silencing LC to dCA1 cells. In contrast, mice with opto-D1 activation in excitatory neurons showed normal contextual memory linking (Fig 7D bottom). This result demonstrates that restoring D1 receptor signaling to excitatory neurons in dCA1 is sufficient to rescue the impairment in contextual memory linking caused by inhibition of LC neuromodulation in dCA1.

Together, these findings demonstrated that LC neurons projecting to dCA1 modulate contextual memory linking by regulating the co-allocation of contextual memories to dCA1 neuronal ensembles in a dopamine-dependent manner. This discovery of a neuromodulatory system that specifically affects memory linking without affecting memory formation, reveals a fundamental separation between the brain mechanisms that modulate these two distinct processes.

## Discussion

The results presented here show that LC neurons projecting to dCA1 modulate the ability of one contextual memory to be linked or associated with another across time, so that the recall of one memory triggers the recall of the other. In the absence of dopaminergic neuromodulation from LC neurons projecting to dCA1, mice show normal contextual memory, but they are unable to link two contextual memories acquired 5 hours apart. These results demonstrate the involvement of the LC in modulating contextual memory liking processes in dCA1, and they reveal that the circuit mechanisms that modulate memory linking can be separate from those that modulate individual memories.

Previous studies have shown that the dorsal hippocampus is critical for contextual memories [78–81]. Studies suggested that dCA1 and dCA3 are involved in different aspects of contextual/spatial memory processing [81–84]. While CA1 represented objects with shared episodes as more similar than those in different episodes, CA3 differentiated the objects encountered in the same episode better [84]. Computational models have proposed that the CA3 region primarily processes information within single episodes, while the CA1 region acts primarily as a comparator that extracts and integrates information across episodes [84–87]. CA1 was shown to process information about temporally ordered sequences of events with the neural code slowly changing over time, while the CA3 population activity remains highly stable [88–91], allowing these two regions to manage different behavioral functions in extended time scales.

These differences between CA1 and CA3 led to the possibility that they might be differentially regulated by neuromodulation. Indeed, our results confirmed a recent study showing that the LC terminals projecting to dCA3, but not those projecting to dCA1, are necessary for single-trial contextual memory formation [34]. Remarkably, we found that the LC cells projecting to dCA1 modulate the ability of one contextual memory to be linked to another acquired close in time (i.e., 5 hours apart). Attesting to the singular role of this projection, another neuromodulatory input to the dCA1, the RN, was not required for contextual memory linking. These results provide compelling evidence for a long-range and specific neuromodulatory pathway that regulates two distinct cognitive functions by modulating two separate subfields of the same brain structure: while dCA3 neuromodulation is critical for contextual memory formation, dCA1 neuromodulation is essential for contextual memory linking.

### Dopaminergic receptor function in dCA1 is critical for contextual memory linking

We further showed that a dopamine D1/D5 antagonist infused bilaterally to the dCA1 disrupted contextual memory linking, while inhibition of β-adrenergic receptors did not. Additionally, we show that optogenetic activation of D1 receptor function in a subset of dCA1 cells can rescue the impairment of contextual memory linking triggered by chemogenetic inhibition of LC cells projecting to dCA1. These findings indicate that the LC cells projecting to dCA1 modulate memory linking in a dopamine-dependent manner.

The focus of LC studies has been traditionally on noradrenergic function (reviewed in [23]). Recently, however, the LC has been shown to co-release dopamine [23–28]. Dopamine is a precursor of noradrenaline, synthesized by tyrosine hydroxylase (TH), not only in dopaminergic, but also in noradrenergic neurons, and is converted by dopamine-β-hydroxylase to noradrenaline in synaptic vesicles [92, 93]. Previous studies suggested that the dopamine to noradrenaline conversion is not complete, and that residual dopamine can be co-released with noradrenaline in noradrenergic terminals, a result that was more recently confirmed using optogenetics and high-performance liquid chromatography in the dorsal hippocampus [25], as well as with pharmacology and microdialysis in the medial prefrontal and occipital cortices [92–95]. Our results showed that contextual memory linking was impaired by inhibition of dopamine D1/D5 receptors, and that it was not affected by inhibiting β-adrenergic receptor function in dCA1. This finding is consistent with previous studies that revealed a role for dopamine released from the LC in memory [25, 26] [34] and LTP [96].

### LC neurons projecting to dCA1 regulate memory linking by modulating neuronal excitability and the properties of memory assemblies

Finally, we show that LC neurons projecting to dCA1 regulate the persistence of increases in neuronal excitability after context exploration, thus biasing neuronal allocation of a subsequent memory to the same ensemble that encoded the first memory. Learning is known to induce a transient increase in neuronal excitability [44–49], creating a temporary window during which future and recent memories can be co-allocated to overlapping populations of neurons [1, 3–5, 50–52]. Manipulation of this excitability by optogenetic or chemogenetic alterations has been shown to disrupt the co-allocation of neuronal ensembles, and impair linking of memories across time [1, 3]. Our results describe a neuromodulatory circuit where LC neurons projecting to dCA1 are critical in maintaining this window of increased excitability triggered by context exploration. Further, this increase in excitability leads to the co-allocation of memories acquired close in time to overlapping dCA1 ensembles, as shown by our *in vivo* and *in silico* findings that inhibition of LC neurons projecting to dCA1 decreases the overlap between these neuronal ensembles. Although inhibition of LC cells projecting to dCA1 did not impact the total number of dCA1 cells activated by learning, as shown with calcium imaging and cFos detection as well as recapitulated in our model, this manipulation impaired how those cells would process subsequent information. Similarly, studies have shown that optogenetic activation of LC projections to dCA1 did not affect the acute activation of dCA1 cells, although it could promote place cell reorganization around a reward site [97].

We also found that the stability of dCA1 coactivity patterns is impaired by the inhibition of LC cells projecting to dCA1 [98]. Importantly, this finding was confirmed by both general pairwise correlations and by methods of cell assembly detection that identify groups of neurons that fire together. Recent findings suggested that the short-term (minutes to hours) stabilization of neuronal assemblies is mediated by changes in intrinsic excitability [98]. Moreover, dopaminergic modulation of dCA1 leads to increases in neuronal coactivity, post learning reactivation of these coactivity patterns, and improved behavioral performance [99]. This is in line with evidence showing that coactivity patterns in dCA1 encode spatial, contextual, and salient features of experience that shape future behavior [56, 57, 100]. The temporal correlation between neuronal activity in dCA1encodes more information than individual neurons, and becomes stronger with learning, slowly building stable networks that correlate with behavioral performance [101]. Interestingly, disrupting the temporal correlation between dCA1 neurons, but not between dentate gyrus neurons, significantly impairs spatial decoding [102].

The LC is known to modulate neuronal output in the dCA1, regulating synaptic plasticity as well as intrinsic excitability through both its dopaminergic and noradrenergic projections [26, 103, 104]. Numerous pharmacological studies have supported the role of D_1_/D_5_ receptors as a gating mechanism for the persistence of plasticity in the hippocampus [96, 105–110]. For example, optogenetic activation of TH+ neurons in the LC, and related D1/D5 receptor activation in CA1, was shown to support long-lasting synaptic potentiation, as well as memory enhancement [26]. Our results now demonstrate that LC activation underlies the persistence of neuronal excitability required for the overlap between dCA1 memory ensembles underlying contextual memory linking. Since we found that D1/D5 receptor activation is required for contextual memory linking, it is possible that the LC modulates neuronal excitability in the dCA1 through the activation of D1/D5 receptors.

All together the results presented here demonstrated that LC cells projecting to dCA1 have a key permissive role in contextual memory linking, without affecting contextual memory formation, and that this effect is mediated by dopamine and not by noradrenaline. Additionally, our results show that these effects on memory linking are caused by the role of these LC cells in modulating increases in excitability of dCA1 neurons triggered by contextual learning, and therefore the co-allocation of contextual memories to overlapping neuronal ensembles in dCA1, as well as the stability of activity patterns within these ensembles. Source and relational memory problems are often associated with a number of neuropsychiatric conditions, including schizophrenia and major depression [7–10]. A key component of these complex cognitive problems is the inability to properly connect and link information about items and events acquired at different times. Therefore, this study sheds light into the mechanisms underlying these deficits, and opens the door to the development of new treatments.

## Author contributions

AC, AL, GF, DA and AJS contributed to the study design. AC designed, performed and analyzed the LC experiments with support from AL, ER and MT. AL designed, performed and analyzed RN experiments. AL performed the detailed analyses of the miniscope experiments with support from DA and AC. GF designed, performed and analyzed the electrophysiological experiments with support from AC and AL. DA and AL wrote the code used for miniscope analyses. AT, GK and PP designed, implemented and analyzed the neurocomputational model. AC and AJS conceptualized the study and wrote the paper together with all the other authors.

## Acknowledgements

We thank Yang Shen, Megha Sehgal, Ying Cai and Andre Sousa for technical support and advice. This work was supported by grants from the NIMH (R01 MH113071), NIA (R01 AG013622), NINDS (RO1 NS106969), from the Dr. Miriam and Sheldon G. Adelson Medical Research Foundation to A.J.S and the Swiss National Science Foundation (SNSF), postdoc mobility fellowship to AC. The computational modelling work was supported by the European Commission (H2020-FETOPEN-2018-2019-2020-01, FET-Open Challenging Current Thinking, GA-863245), the NIH (R01MH124867-01) and the Einstein Foundation Berlin.

## Methods

### Animals

10-12 week-old male and female C57BL/6NTac mice were purchased from Taconic Farms (Germantown, NY) for all experiments. Mice were group housed with free access to food and water, and maintained on a 12:12 hour light: dark cycle. Two weeks before an experiment, they were single-housed. All experiments were performed during the light phase of the cycle. All studies were approved by the Chancellor’s Animal Research Committee at UCLA.

### Immunostaining

Mice were transcardially perfused with 4% PFA (4% paraformaldehyde in 0.1 M phosphate buffer) and after perfusion, brains were kept in the fixation solution overnight at 4 °C, then transferred to 30% sucrose solution for 24 h, sectioned (40 μm thickness) on a cryostat and stained while free-floating.

The sections were blocked for 1 h at room temperature in 0.5% Triton-X100 in PBS (PBST) and 10% normal goat serum (Vector Laboratories, S-1000) solution. The subsequent primary and secondary antibodies were diluted in the same blocking solution. The primary antibody incubation was overnight (~24-36 h) at 4 °C, and the secondary antibody incubation was 2 h at room temperature, both with constant shaking.

Primary antibodies: chicken anti-GFP (Abcam AB13970, 1:1000), rabbit anti-cFos (Cell Signaling, 9F6, #2250, 1:500), chicken anti-TH (Abcam AB76442), guinea pig anti-RFP (SySy 390 004), rabbit anti-RFP (Rockland antibodies 600-401-379), rabbit anti-serotonin transporter antibody (Millipore Sigma, AB9726, 1:500) were used for immunostaining. Brain slices were incubated with 4’,6-diaminodino-2-phenylindole (DAPI, Invitrogen, 1:1000) for 15 min, washed with PBST two times and PBS once before mounting onto slides.

Secondary antibodies were Alexa Fluor 488, 568 and 647 (Invitrogen).

Immunostaining images were acquired with a Nikon A1 Laser Scanning Confocal Microscope (LSCM) and analyzed with automatic spot-detection algorithm (Imaris 9.2, Bitplane AG).

### Viral constructs

CAV2-cre and CAV2-Flp were purchased from Plateforme de Vectorologie de Montpellier, IGMM, France [40, 41].

pAAV-hSyn-DIO-hM4D(Gi)-mCherry was a gift from Bryan Roth (Addgene viral prep # 44362-AAV8; http://n2t.net/addgene:44362 ; RRID:Addgene_44362) [42]. pAAV-hSyn-DIO-mCherry was a gift from Bryan Roth (Addgene viral prep # 50459-AAV8;http://n2t.net/addgene:50459;RRID:Addgene_50459).

AAV.Syn.GCaMP6f.WPRE.SV40 was a gift from Douglas Kim & GENIE Project (Addgene viral prep # 100837-AAV1; http://n2t.net/addgene:100837; RRID: Addgene_100837) [111] pAAV-hSyn-DIO-EGFP was a gift from Bryan Roth (Addgene viral prep # 50457-AAV8; http://n2t.net/addgene:50457; RRID:Addgene_50457),

AAV-8-Ef1a-fDIO DREADD Gi-mCherry (GVVC-AAV-171) and AAV-8-Ef1a-fDIO-mCherry-WPRE (GVVC-AAV-155) were purchased from Neuroscience Gene Vector and Virus Core at Stanford, pAAV-Ef1a-fDIO EYFP was a gift from Karl Deisseroth (Addgene viral prep # 55641-AAV1; http://n2t.net/addgene:55641 ; RRID:Addgene_55641) [71]. pAAV.hSyn.Cre.WPRE.hGH was a gift from James M. Wilson (Addgene viral prep # 105553-AAV1, http://n2t.net/addgene:105553 ; RRID:Addgene_105553). pENN.AAV.CamKII 0.4.Cre.SV40 was a gift from James M. Wilson (Addgene viral prep # 105558-AAV1; http://n2t.net/addgene:105558; RRID:Addgene_105558).

AAV5-EF1a-DIO-OptoD1-EYFP (UNC, Chapel Hill) [72].

### Stereotaxic Surgery

Animals were anesthetized with 3-4% isoflurane and maintained at 1.5-2% in a stereotaxic head frame on a heat pad. Artificial tears were applied to the eyes to prevent eye drying. A midline incision was made down the scalp, and a craniotomy was performed with a dental drill. After surgery, the animals were subcutaneously injected with Carprofen (5 mg/kg) and Dexamethasone (0.2 mg/kg) before recovery. Water with amoxicillin was provided for two weeks.

For virus injection, a Nanoliter injector (World Precision Instruments) was used to infuse virus with Micro4 Controller (World Precision Instruments), injecting at coordinates relative to bregma/skull (mm): dCA1 at −1.8 (AP), +/− 1.5 (ML), −1.6 (DV); dCA3 at −1.8 (AP), +/− 2.0 (ML) and −2.0 (DV); LC at −5.45 (AP), +/− 1.2 (ML), −3.65 (DV), RN at 4 mm (AP), −1.2 (ML at 15 degree angle), −4.5 (DV). Virus was infused at 50nL/min. After infusion, the capillary was kept at the injection site for 10 min and then withdrawn slowly. The incision was sutured and the mice were allowed to recover for 2 weeks before start of behavior.

For cannula implantation, two guide cannulas (Plastics One, C313GS-5/SPC) were implanted at the following coordinates relative to bregma (mm): AP: −2.1, ML: ±1.7 for dCA1. Three weeks after cannulation, mice were anesthetized and sterilized saline or drug (doses as mentioned in text, 300nl, 100nL/min) was infused into hippocampus through the internal cannula (Plastics One, C313IS-5/SPC) at DV:-1.65 relative to skull. After infusion, the internal cannula was left in place for an additional 5 min to ensure full diffusion.

For optical fiber implantation, fiber Optic Cannula (Newdoon, 200 μm, NA=0.37) was immediately implanted after virus injection. The tip of the optic fiber was placed 1mm above the virus injection site. Then, the canula was fixed with Metabond and dental cement.

For miniscope implantation, a GRIN lens was implanted into the dorsal hippocampal CA1 region as previously described [1]. After GCaMP6f virus injection, a ~2mm diameter circular craniotomy was centered at the injection site. The cortex directly below the craniotomy was aspirated with a 27-gauge blunt syringe needle attached to a vacuum pump. Cortex buffer (NaCl 135mM, KCL 5mM, CaCl2 2.5mM, MgSO4 1.3mM, HEPES 5mM, PH 7.4) was repeatedly applied to the exposed tissue to prevent drying. The GRIN lens (0.52 NA, 1.8 mm in diameter, Edmund Optics) was slowly lowered above CA1 to a depth of 1.35 mm ventral to the surface of the skull at the most posterior point of the craniotomy. Next, a skull screw was used to anchor the lens to the skull. Both the lens and skull screw were fixed with super glue (Loctite, 45198) and dental cement (Jet Denture Repair Package, Lang, 1223CLR). Low Toxicity Silicone Adhesive (Kwik-Sil, World Precision Instruments) was used to cover the GRIN Lens for protection. Two weeks later, a small baseplate was cemented onto the animal’s head atop the previously formed dental cement.

### Behavioral procedures

Two weeks after surgeries, the mice were first handled for 5 days (2min/day) in their housing room, and then habituated to transportation and external environmental cues for 2 minutes in the experimental room each day where they were also handled (2min/day) for another 3 days. In the contextual memory linking task, mice explored 2 different contexts (A and then B) which were separated by 5 hours. Mice explored each context for 10 min. CNO injection (5mg/kg, i.p) or drug infusions were done as noted. For immediate shock, two days later, mice were placed in chamber B for 10 s followed by a 2s shock (0.75 mA). 58 seconds after the shock, mice were placed back in their home cage. For the context tests, mice were returned to the designated context for the next three days (A, B and a new neutral context C) in a counterbalanced manner. Contexts A and C were counterbalanced. Freezing was assessed via an automated scoring system (Med Associates) with 30 frames per second sampling; the mice needed to freeze continuously for at least one second before freezing could be counted.

For single contextual memory, the mice were handled and habituated the same way. They explored one context for 10min, and two days later, the mice were placed in that chamber for 10 s followed by a 2s shock (0.75 mA). 58 seconds after the shock, mice were placed back in their home cages. For the context tests, mice were returned to the shocked context and a counterbalanced neutral context.

### Optogenetics

Two weeks after virus injection and optic cannula implantation, the mice were handled for 5 days (2min/day) in their housing room, and then handled in their experimental room for another 5 days where they were additionally habituated with the optic fiber connected in their home cage (2min/day). On the day of behavioral linking, they were systemically (i.p) injected with CNO (5mg/kg) 30min before context A where they received light stimulation during the 10 min exploration (473nm, 8-10mW, 50s on/10s off). Five hours later, they were taken to explore context B for 10min without drug or optic fiber.

### Computational modeling

A previously published model network of memory allocation was adapted [64]. The model network consists of populations of excitatory and inhibitory neurons which are modeled as 2-stage integrators to account for dendrites. Neurons consist of a somatic unit connected to independent dendritic subunits. Both dendrites and soma are modeled using simplified integrate-and-fire model dynamics, where the somatic unit includes adaptation current, while dendritic units receive excitatory synaptic currents. Dendrites and soma are coupled via axial resistance. Inhibitory neurons provide feedback inhibition and are separated in 2 equal sub-populations, soma-targeting and dendrite-targeting. Each dendritic subunit integrates incoming the synaptic inputs which reside on it independently as follows:

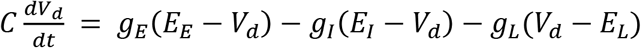

Where *V*_*d*_ is the dendritic depolarization, *C* is the membrane capacitance of, *E*_*E*_ is the reversal potential for excitatory receptors, *E*_*I*_ is the reversal potential for inhibitory receptors, *E*_*L*_ is the resting potential (0mV), *g*_*L*_ is the leak conductance and *g*_*E*_, *g*_*I*_ are the instantaneous activations of synaptic currents

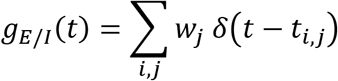

Where *w*_*j*_ is the weight of synapse *j* and *t*_*i,j*_ are the timings of incoming spikes.

Somatic spiking follows an Integrate and Fire model with adaptation:

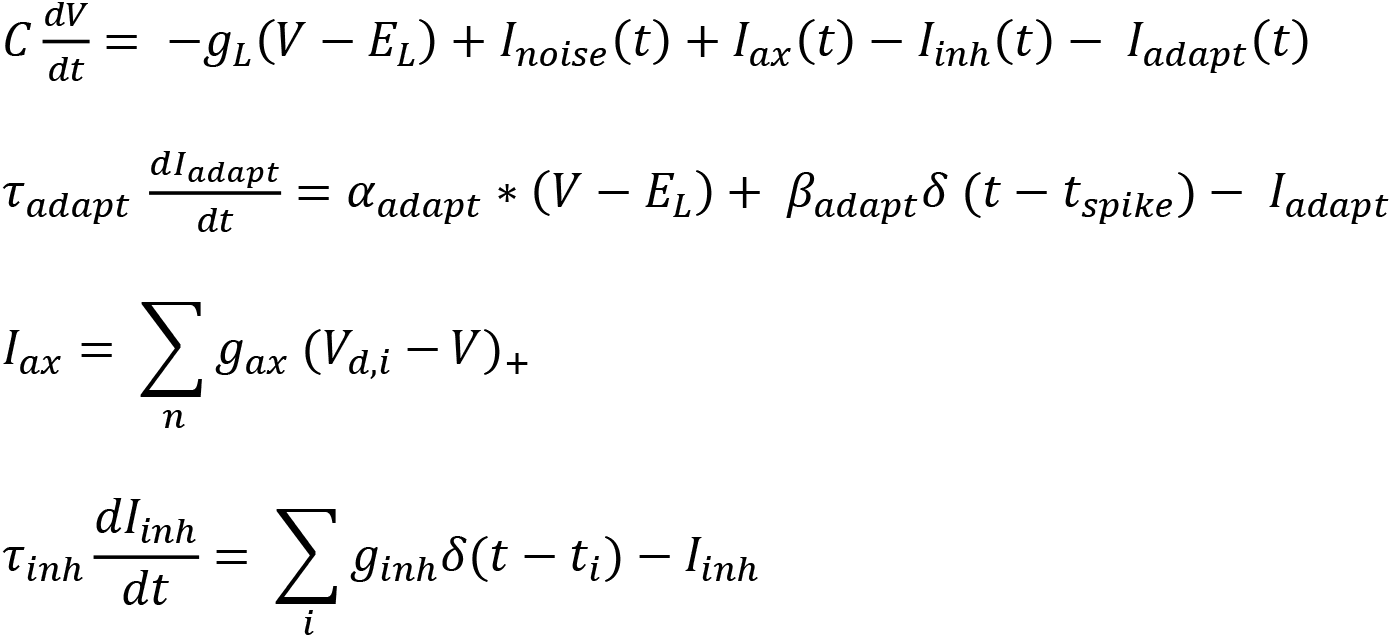

Where *V* is the somatic voltage, *I*_*noise*_ is uniform noise current (max amplitude 500 pA), *I*_*ax*_ is the excitatory axial current, *I*_*inh*_ is the filtered inhibitory input from somatically-targeting interneurons, *I*_*adapt*_ is the adaptation current, *τ*_*adapt*_ is the adaptation time constant, *α*_*adapt*_ the adaptation coupling parameter, *β*_*adapt*_ is the amount by which adaptation current increases every time the neuron spikes, *g*_*ax*_ is the axial resistance, *τ*_*inh*_ is the time constant of inhibitory current and *g*_*inh*_ the inhibitory current scaling constant.

Somatic spiking occurs when the somatic voltage reaches the spike threshold *θ*_*soma*_. Calcium influx is recorded on the level of single synapses, branches and whole neuron. Synaptic and dendritic branch calcium is increased when a presynaptic spike coincides with back propagating action potential. Somatic calcium is increased by every time a somatic spike occurs. For plasticity-induction purposes the overall sum of accumulated calcium at the end of a 4 second stimulation is used.

Synapses representing memories to be encoded are initially allocated randomly to the dendritic subunits of pyramidal neurons with initial weight 0.4. In addition, feedback synapses between pyramidal and inhibitory populations are allocated at random, with separate distributions for soma-targeting and dendrite-targeting interneurons.

Calcium influx in a synapse during a 4 second stimulus presentation determines plasticity, which is dependent on the availability of plasticity-related proteins (PRPs). Synapses are selected for plasticity according to a Hebbian rule: Synapses that receive significant calcium influx and reside on a neuron that is highly activated are selected for potentiation, otherwise they are selected for depression. The update of the weights of the selected is dependent on the level of Plasticity-Related-Protein (PRP) synthesis, which is assumed to be somatic. The level of PRPs after the somatic calcium level exceeds the threshold *Θ*_*PRP*_ over time in minutes follows the alpha function

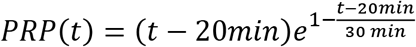

The sum of available proteins determines the rate of consolidation of the weights *w* of synapses. Synaptic weights are also subject to a homeostatic plasticity rule, which normalizes the total synaptic input to each neuron over long time scales:

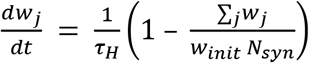

Where *w*_*init*_ is the 0.3, *N*_*syn*_ the total number of synapses in the neuron and *τ*_*H*_ the time constant of homeostatic synaptic scaling.

Simulation of memory formation takes place according to the following protocol: For every memory being encoded, the inputs which represent the memory are stimulated for 4 seconds with a high firing rate (80Hz) to drive the firing of the initially-weak synapses. After the first memory encoding, protein dynamics and excitability modulation are modeled for 5 hours as detailed in [64] and then the second memory is encoded. The neurons which have firing rate >10Hz during encoding are considered as highly-active and thus part of the memory engrams. Under control conditions, the excitability of these neurons is increased after encoding for a period of up to 12 hours. Under the LC block condition, this increase in excitability does not take place. The overlap between active populations is measured by the ratio of (neurons active in both memories) / (neurons active in either memory). For additional details of the model see [64]. The parameters used are listed in Supplementary Table 2. The model was written in C++. The source code for the simulation, data analysis and scripts to reproduce the data and figures are available in the ModelDB database (Accession Number 267173).

### Slice Preparation

Adult mice injected with a CAV-Cre virus in the dCA1 and either AAV-hSyn-DIO-hM4D(Gi)-mCherry or AAV-hSyn-DIO-mCherry in the LC were administered clozapine-N-oxide (CNO, 5mg/kg, i.p) 30 min prior to context exploration. The mice were allowed to explore a novel context for 10 min and five hours later were deeply anaesthetized with isoflurane and decapitated. The brain was rapidly dissected out and transferred to oxygenated (95% O_2_ / 5% CO_2_), ice-cold cutting solution containing (in mM): 93 NMDG, 93 HCl, 2.5 KCl, 1.2 NaH_2_PO_4_, 30 NaHCO_3_, 20 HEPES, 25 glucose, 2 Thiourea, 5 Na-ascorbate, 3 Na-pyruvate, 5 N-acetyl-L-cysteine, 2 CaCl_2_ and 2 MgCl_2_. Coronal slices (300 μm thick) containing the hippocampus were cut using a Leica VT1200 vibrating blade microtome, transferred to a submerged holding chamber containing oxygenated cutting solution and allowed to recover for 15 min at 34°C. Following recovery, the slices were transferred to an oxygenated solution containing (in mM): 92 HEPES, 2.5 KCl, 1.2 NaH_2_PO_4_, 30 NaHCO_3_, 20 HEPES, 25 glucose, 2 Thiourea, 5 Na-ascorbate, 3 Na-pyruvate, 5 N-acetyl-L-cysteine, 2 CaCl_2_ and 2 MgCl_2_ and allowed to recover further for 1hr. Following incubation, slices were transferred to a superfused recording chamber and constantly perfused with oxygenated aCSF containing (in mM): 115 NaCl, 10 glucose, 25.5 NaHCO_3_, 1.05 NaH_2_PO_4_, 3.3 KCl, 2 CaCl_2_ and 1 MgCl_2_ and maintained at 28°C.

### Whole-cell patch recordings

Whole cell current-clamp recordings were performed on pyramidal neurons in the dorsal CA1 region of the hippocampus using pipettes (3-5MΩ resistance) pulled from thin-walled Borosilicate glass using a Sutter P97 Flaming/Brown micropipette puller and filled with an internal solution containing (in mM) 120 K-methylsuphate, 10 KCl, 10 HEPES, 10 Na-phosphocreatine, 4 Mg-ATP and 0.4 Na-GTP. All recordings were obtained using a MultiClamp 700B amplifier controlled by the pClamp 10 software and digitized using the Digidata 1440A system. Signals were filtered at 10kHz and digitized at 20kHz. Neurons were included in the study only if the initial resting membrane potential (Vm) ≤ −55 mV, access resistance (Ra) was <25MΩ and were rejected if the Ra changed by >20% of its initial value. For all recordings, neurons were held at −65 mV. The stable resting membrane potential of neurons were measured and averaged over a 60s duration with 0mA current injection immediately after breaking in. Input resistance was measured as the slope of the steady-state voltage response to increasing current injections (−50pA to 50pA, Δ=10pA). To investigate the firing rate of neurons, the number of action potentials fired in response to a 600 ms pulse of depolarizing current injection (0pA to 480pA in 20pA increments) was calculated. Three pulses were delivered for each current amplitude and the average number of action potentials fired for each current amplitude was plotted. The recordings were analyzed using Stimfit 0.15.8.

### Miniscope analysis

One-photon calcium imaging was recorded using UCLA miniscopes [1, 112]. During recordings, digital imaging data were sent from the CMOS imaging sensor (Aptina, MT9V032) to custom data acquisition (DAQ) electronics and USB Host Controller (Cypress, CYUSB3013) over a light weight, highly flexible co-axial cable. Images were acquired at 30 frames per second, using display resolution at 752 x 480 pixels (1 pixel = 1-2μm), and saved into uncompressed avi files. The analysis pipeline was written in MATLAB using first the NoRMCorre algorithm for motion correction (rigid registration) [113], followed by individual neuron identification and extraction using the CNMF-E algorithm [114]. During motion correction, videos were 2x spatially downsampled using the default built-in NoRMCorre protocol. During CNMF-E initialization, videos were further 2x spatially down-sampled and 5x temporally down-sampled. After motion correction, the videos were analyzed in two different ways. In the single session analysis, videos from individual sessions were directly input for CNMF-E processing; while in the concatenated analysis, these videos were first aligned and then concatenated before CNMF-E analysis (Concat Pipeline). Alignment was performed using a semi-automatic alignment tool based on the “imregtform” function (Matlab – image processing toolbox) followed by manual checking of landmarks (usually blood vessels).

After concatenation and CNMF-E processing, the quality of neuron extraction in the concatenated analysis was verified using a MATLAB custom-made Neuron Deletion GUI. Putative false-positive neurons were filtered out using the following exclusion criteria: 1) abnormalities on ROI morphology or calcium trace, and 2) calcium trace peaks with no corresponding fluorescence increases in the video. Experimenters were blinded to all steps of the analysis. All downstream analysis was performed using the remaining ROIs after filtering (putative neurons). Neurons detected on the CNMF-E analysis on single 10-min sessions were only used if they found a correspondent match on the filtered concatenated analysis. We used the spatial foot prints (neuron.A, output from CNMF-E) from each one of the detected cells for the binary matching analysis between each one of the single sessions and the filtered concatenated analysis. The centroid distance and spatial correlation were calculated for all cell pairs (concatenated x single session). Cells were deemed as a match if their spatial correlation ≥ 0.8 and their centroid distance ≤ 5 pixels. We defined the percentage overlap between 2 given sessions (e.g., A and B) as the ratio between the intersection (*A*⋂*B*) and the union (*A*⋃*B*) among the cells activated in the respective sessions, as in the formula:

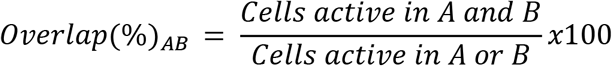

The probabilistic calculations on activated cells were defined as the conditional probability of cells to have been retrospectively active during LC manipulation 5h in the past, given that they are active during exploration of the following context. This indicates the likelihood that the cells activated during exploration of a novel context were biased by the cells activated during exploration of another novel context 5h before, and is formally calculated as:

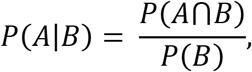

where *P*(*A*⋂*B*) and *P*(*B*) are respectively the probability of cells being active in A and B and the probability of cells being active in B. These probabilities were defined as the ratios *P*(*A*⋂*B*) = *Cells active in A and B/U* and *P*(*B*) = *Cells active in B*/*U*, in which *U* was defined as the universe (total number) of cells detected in the concatenated analysis.

For the analysis of neuronal activity dynamics across sessions, the raw calcium traces (neuron.C_raw) extracted from putative neurons using CNMF-E were deconvolved into spike probabilities using the foopsi thresholded method (OASIS toolbox) [115]. Finally, the spike probabilities from single frames were binarized between 1 (active) and 0 (inactive). Mean firing rate was calculated as the number of active frames per second within the overlapping neuronal population in each context. The coactivity map from a specific session was defined as the matrix containing all the pairwise temporal correlations (PWC, Pearson Correlation) between the binned activity (100 ms) of any two given neurons. Mean PWC was calculated as the average of all values within a coactivity map of overlapping neurons, discarding autocorrelations (correlation between a given neuron and itself).

PWC stability represents the stability of how neurons fire together across both contexts. PWC stability is calculated by converting the coactivity values for all overlapping neurons with themselves in each session into a vector, and then measuring the Pearson linear correlation between these vectors. To avoid potential high-correlation false positives due to fluorescence leak through between putative neurons that are spatially close to each other, we excluded the correlation-pairs between neurons that had overlapping spatial footprints or had centroids closer than 20 pixels from one another. PWC stability values closer to 1 means that neurons tend to keep their partnership profile (fire together, show independent activity, or fire when the other remains silent) in both contexts. Representative graphs to visualize the PWC stability (Fig. 4E) were generated by calculating the absolute delta between the PWC matrices in context B vs context A. To prevent ensemble size effects, PWC stability was calculated in subsamples of the neurons from individual animals (1000 permutations), equal to the smallest number of overlapping neurons detected in all animals. The final PWC stability value for each animal was defined as the average of these 1000 values.

Cell assemblies within the overlapping neuron population were identified using a PCA/ICA mathematical tool based on Hebbian co-firing rules [63]. This method identifies a number of co-activation patterns for every dataset (in the case of our dataset, between 1-6 patterns per session for the overlapping neuron population) characterized by a linear correlation between the activity of the cells within each assembly. For each assembly detected, every recorded cell will have a weight representing how much that cell fires together with other cells that participate in the assembly. This weight score can go from +1 to −1, corresponding to a range of perfect correlation to perfect anti-correlation between the firing of that specific neuron and the activity of the assembly pattern. A high absolute weight means the neuron is a part of that assembly, while a low absolute weight means it is not a part of the assembly.

The cell assembly detection framework can be summarized by these three main steps:

1. Generation of the Activity Matrix: the neuronal activity was binned (100ms) and normalized by z-score (variance is set to 1 and mean is set to 0).
2. Detection of Cell Assemblies: PCA was performed in the Activity Matrix to identify its principal components (putative assemblies). Parallel to that, a random circular shift method was used on each neuronal activity independently, breaking any real temporal correlation between neurons. The shifted neuronal activity was used to estimate surrogate Activity Matrices and their respective eigenvalues (200 permutations), which were used to define a random distribution. Principal components of the original Activity Matrix with associated eigenvalues statistically different (95% threshold) from the randomized distribution were considered significant cell assemblies.
3. Generation of Assembly Patterns: Independent Components Analysis (ICA) was performed on the original Activity Matrix projected on the subspace spanned by the patterns of the significant cell assemblies. The output independent components can be understood as assembly patterns in which values attributed to each neuron define the weights of the cells (relative relevance) in the corresponding assembly.

Stability Index (SI) for cell assemblies represents how similar the weights for matching assemblies are maintained across the two contexts. First, a similarity index was calculated by measuring the cosine similarity (inner product) between all identified cell assembly patterns in the overlapping neuron population of context B (second exposure) versus context A (first exposure) [116]. Assemblies of context B were then matched in a 1:1 ratio to the assembly from context A with the highest similarity value. Stability index was then calculated by averaging the similarity values of all matched assemblies.

### Statistical Analyses

The investigators who collected and analyzed the data including behavior, electrophysiological and staining were blinded to the experimental conditions. Error bars in the figures indicate the SEM. All statistical analyses were performed using GraphPad Prism 9.0.2. For behavior and immunohistochemistry experiments, n designates the number of mouse. For electrophysiological measurements, n designates the number of neurons. Statistical significance was assessed by unpaired- t test (two-tailed), Mann-Whitney, one-sample t-test, one-way ANOVA, two-way RM ANOVA where appropriate, followed by the indicated post-hoc tests. Normality was tested by Shapiro-Wilk and Kolmogorov-Smirnov tests. The level of significance was set at p<0.05.

