## Supplementary Material for "A Locus Coeruleus- dorsal CA1 dopaminergic circuit modulates memory linking"

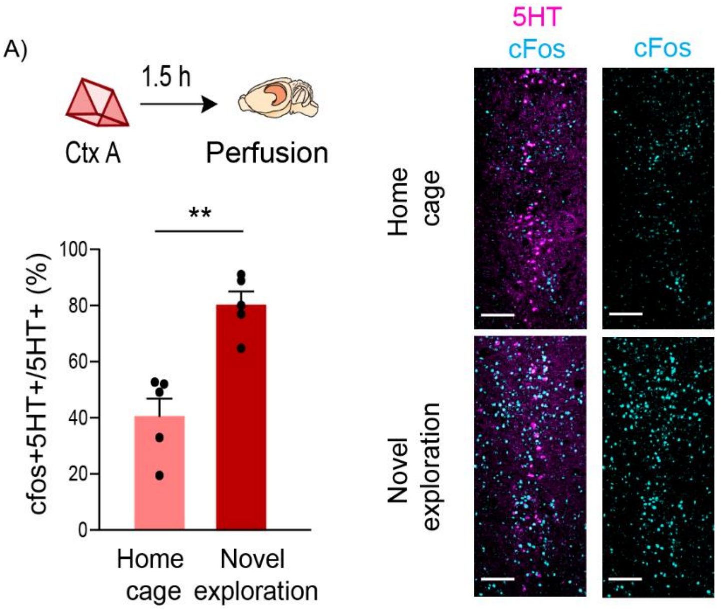


**Suppl. Fig. 1: Context exploration induces cFos expression in RN**

Exploration of a novel context increased cFos expression in 5HT+ positive cells of RN (unpaired t-test, n=5, **p<0.01). Example images of 5HT and cFos staining in RN. Scale bar, 50µm. 5HT- magenta, cFos- cyan.

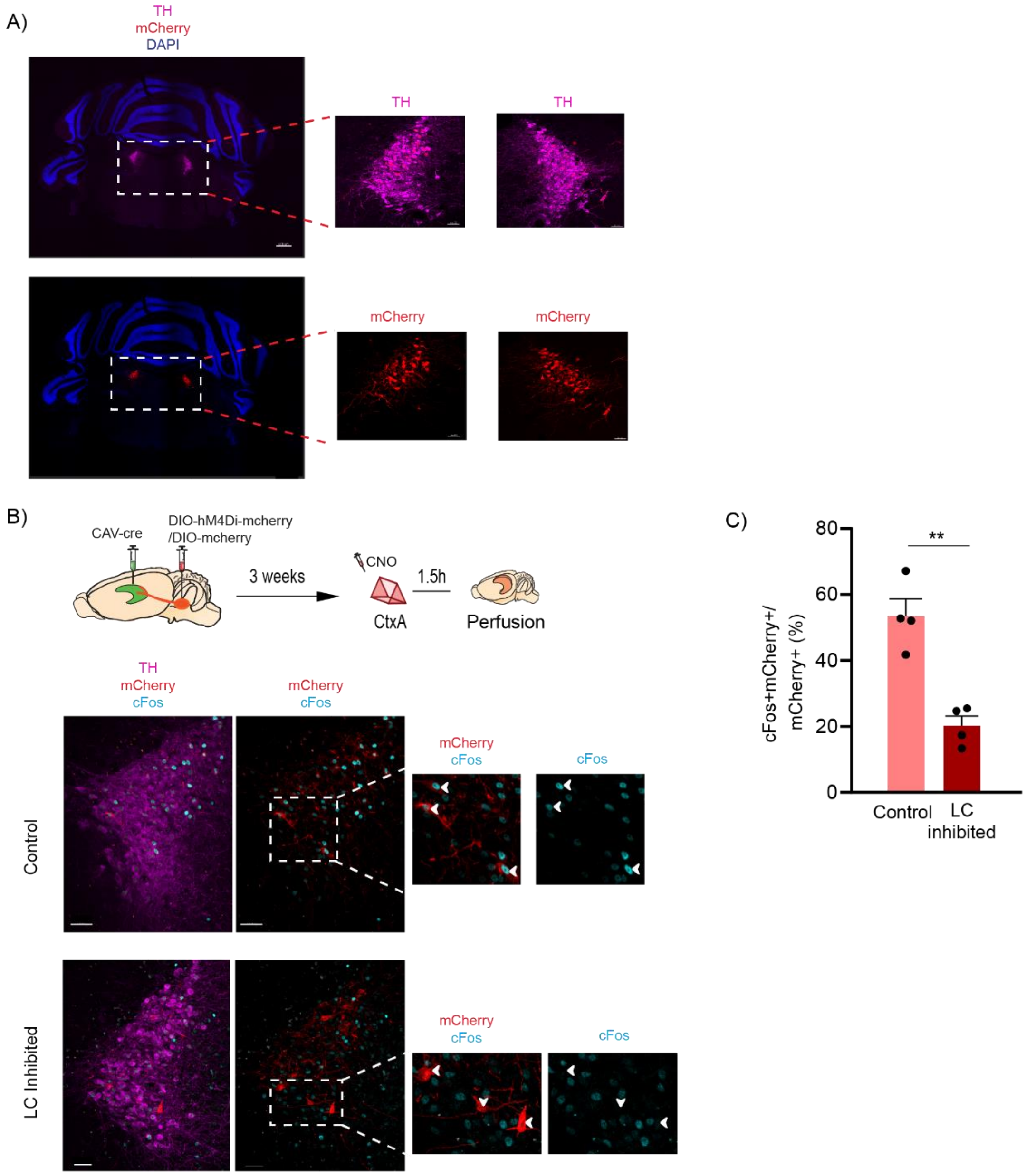


**Suppl. Fig 2: Inhibition of LC cells projecting to dCA1 reduced cFos expression triggered by contextual exploration**

A) Representative images for injection sites at LC to show the spread of DIO-hM4Di-mcherry unfloxed by the CAV-cre retrogradely transported from dCA1. TH- magenta, mCherry- red, DAPI- blue. Scale bar, 300µm.

B) Schematics of the experimental procedure. CAV-cre was injected in CA1 and DIO-hM4Di-mCherry (LC inhibited) /DIO-mCherry (control) in LC. All mice received CNO 30min prior to exploration of context A and were perfused 90min later to check cFos induction in the mcherry-expressing cells in LC. Representative images. TH- magenta, mCherry- red, cFos- cyan. Scale bar, 50µm.

C) Inhibition of LC cells projecting to dCA1 during exploration of a novel context significantly reduced cFos expression in the TH positive cells of LC (unpaired t-test, n=4, **p<0.01).

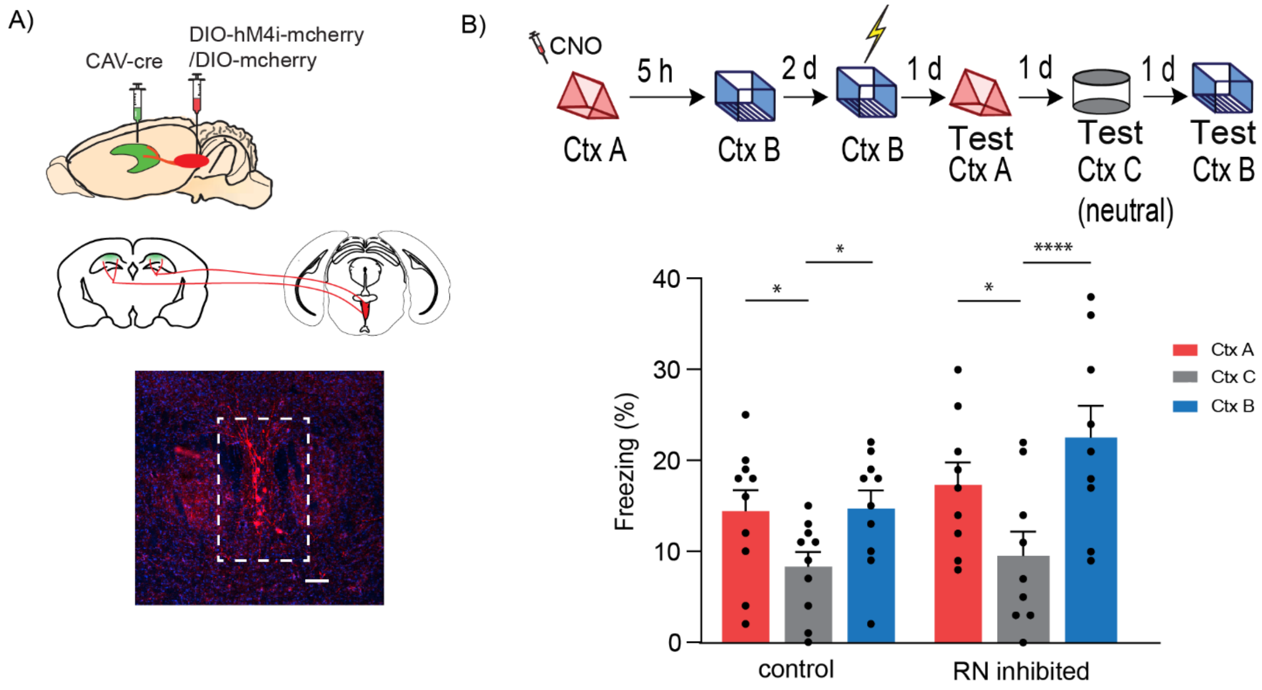


**Suppl. Fig 3: RN to dCA1 projecting cells are not required for contextual memory linking**

A) Schematics of experimental design. CAV-cre was injected in CA1 and DIO-hM4Di-mcherry/DIO-mcherry in RN. Representative images for injection sites at RN to show the spread of DIO-hM4Di-mcherry unfloxed by the CAV-cre retrogradely transported from dCA1. mCherry- red, DAPI- blue. Scale bar, 100µm.

B) Schematics of experimental design. Context A- Ctx A, Context B- Ctx B, Context C- Ctx C (neutral). CNO- Clozapine-N-oxide was given to all mice. Inhibition of RN cells projecting to dCA1 during exploration of context A did not affect the process of memory linking. (Control, n=10; LC inhibited, n=9. Two-way repeated measures ANOVA, Sidak post hoc. $^{*}p < 0.05$, ****$p < 0.0001$)

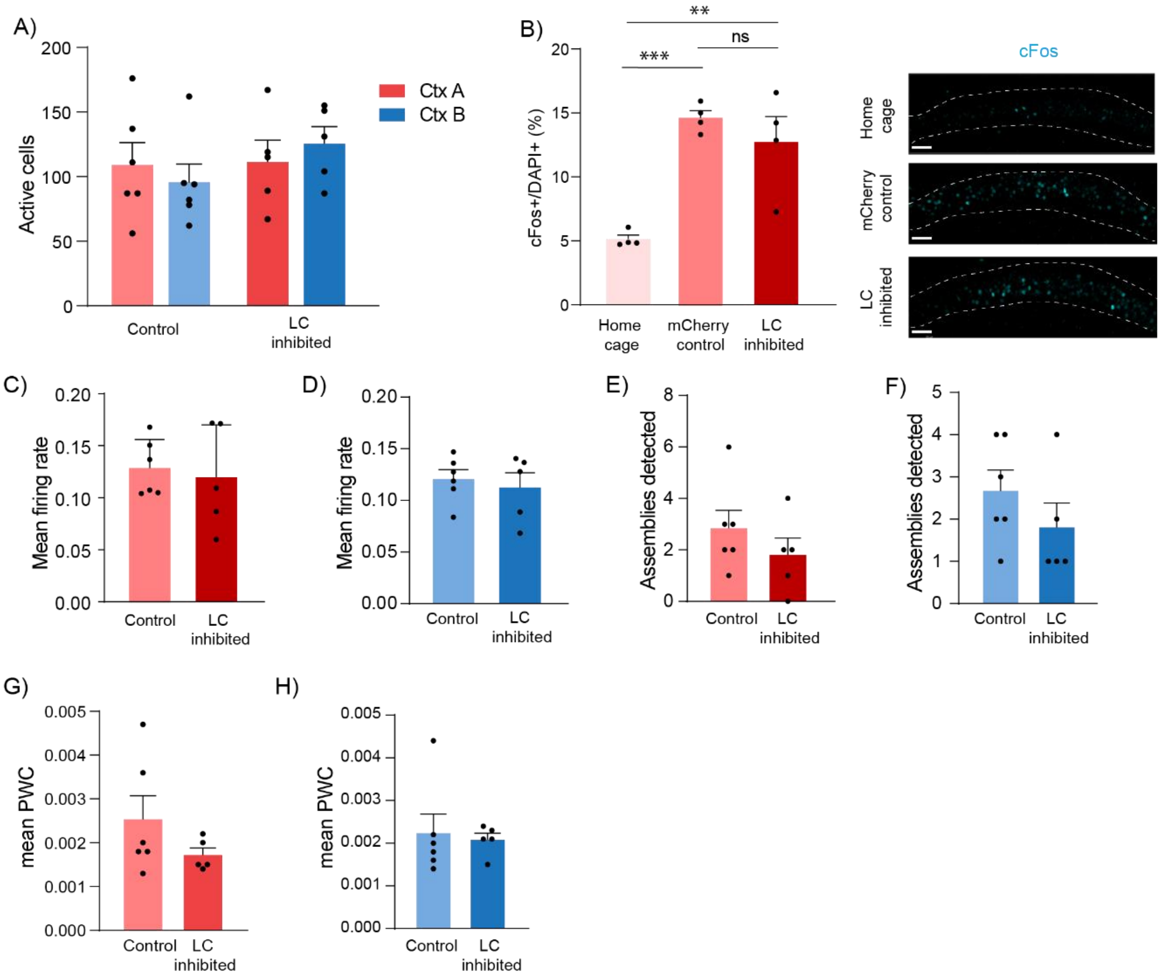


**Suppl. Fig 4: LC to dCA1 inhibition does not affect the properties of individual neuronal ensembles**

A) Total number of active cells detected in dCA1 by calcium imaging does not change in either context upon inhibition of LC cells projecting to dCA1 during exploration of context A. Control n=6, and LC inhibited n=5, Two-way RM ANOVA, Sidak posthoc.

B) Induction of cFos upon novel context exploration in dCA1 was not affected by inhibition of LC cells projecting to dCA1, n= 4 in all groups, One-way ANOVA, Fisher's LSD. Example images for cFos- cyan, principal layer of dCA1 outlined, scale bar, 50 µm.

C) There is no difference in mean firing rates in context A of the neurons active in both contexts A and B (overlapping neurons). Control n=6, and LC inhibited n=5, unpaired t-test.

D) There is no difference in mean firing rates in context B of the overlapping neurons. Control n=6, and LC inhibited n=5, unpaired t-test.

E) There is no difference in total assemblies of the overlapping neurons detected in context A. Control n=6, and LC inhibited n=5, unpaired t-test.

F) There is no difference in total assemblies of the overlapping neurons detected in context B. Control n=6, and LC inhibited n=5, unpaired t-test.

G) There is no difference in mean pair-wise correlation of the overlapping neurons detected in context A. Control n=6, and LC inhibited n=5, unpaired t-test.

H) There is no difference in mean pair-wise correlation of the overlapping neurons detected in context B. Control n=6, and LC inhibited n=5, unpaired t-test.

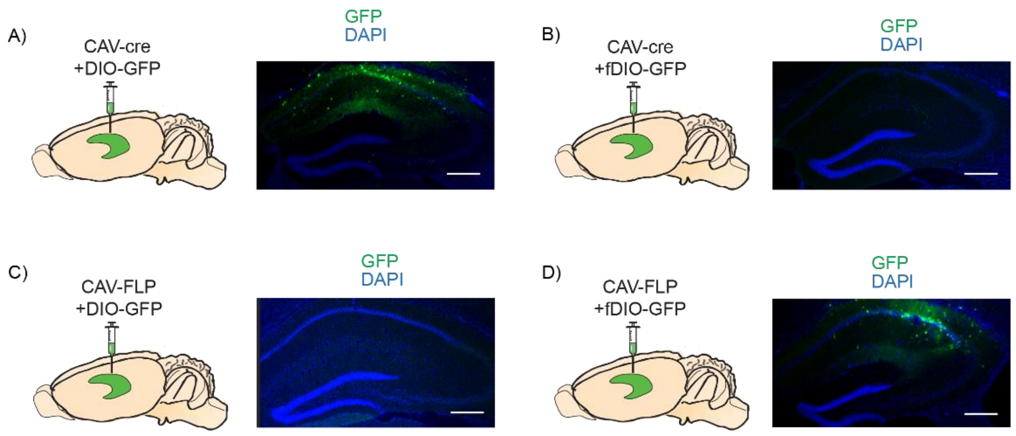


**Suppl. Fig 5: Cre-DIO and Flp-fDIO systems do not interfere with each other**

A) Schematics of experimental design. A cocktail of CAV-cre and DIO-GFP was injected in dCA1. The expression of GFP shows the functionality of the cre-DIO system. GFP- green, DAPI- blue. Scale bar, 50µm.
B) Schematics of experimental design. A cocktail of CAV-cre and fDIO-GFP was injected in dCA1. The lack of expression of GFP confirms that the incompatibility of the two systems. GFP- green, DAPI- blue. Scale bar, 50µm.
C) Schematics of experimental design. A cocktail of CAV-FLP and DIO-GFP was injected in dCA1. The lack of expression of GFP shows confirm that the incompatibility of the two systems. GFP- green, DAPI- blue. Scale bar, 50µm.
D) Schematics of experimental design. A cocktail of CAV-FLP and fDIO-GFP was injected in dCA1. The expression of GFP shows the functionality of the FLP-fDIO system. GFP- green, DAPI- blue. Scale bar, 50µm.

| **Property** | **Control** | **LC inhibited** | **P value** | **Statistical test** |
|---|---|---|---|---|
| RMP (mV) | -70.1 + 0.8 | -71 + 0.5 | 0.36 | Unpaired t-test |
| Rin (MΩ) | 159.7 + 17.1 | 146.3 + 17.1 | 0.55 | Mann Whitney test |

**Suppl. Table 1: Passive membrane properties of dCA1 pyramidal neurons.** There was no difference in the resting membrane potential (RMP) and input resistance (Rin) of neurons from control and LC inhibited slices. Mean ± s.e.m.

| **Parameters of the microcircuit model** | | |
|---|---|---|
| $N_{pyr}$ | Number of excitatory neurons | 400 |
| $N_{inh}$ | Number of inhibitory neurons | 50 dendrite-targeting (DT)<br>50 soma-targeting (ST) |
| $N_{dend}$ | Number of dendritic subunits per neuron | 20 for excitatory<br>1 for interneurons |
| $N_{pyr \rightarrow ST}$ | Total number of synapses from excitatory neurons to soma-targeting(ST) interneurons | 1000 |

| | | |
|---|---|---|
| $N_{pyr \rightarrow DT}$ | Total number of synapses from excitatory neurons to dendrite-targeting (DT) | 1000 |
| $N_{ST \rightarrow pyr}$ | Total number of synapses from ST interneurons to excitatory neurons | 10000 |
| $N_{DT \rightarrow pyr}$ | Total number of synapses from DT interneurons to excitatory neurons | 2000 |
| $N_{input \rightarrow pyr}$ | Total number of weak connections from input afferents to pyramidal dendrites per memory | 5600 |
| $E_L$ | Leakage reversal potential | 0 mV |
| $g_E$ | Dendritic excitatory synaptic conductance | 26 nS |
| $g_I$ | Dendritic inhibitory synaptic conductance | 20nS |
| $g_{Ld}$ , $g_L$ | Dendritic/somatic leak conductance | 50nS |
| $g_{Inh}$ | Somatic inhibitory current scaling constant | 600nS |
| $\tau_{Inh}$ | Somatic inhibitory current time constant | 30msec |
| $E_E$ | Excitatory synapse reversal potential | +70mV |
| EI | Inhibitory synapse reversal potential | -10mV |
| C | Membrane capacitance | 200pF |
| $\tau_{dend}$ | Dendritic membrane time constant | Inhibitory: 20msec<br>Excitatory: 25msec |
| $V_d$ | Dendritic Depolarization | -10mV <Vd< 70mV |
| $g_{ax}$ | Axial conductance | 30nS |
| $\theta_{soma}$ | Voltage threshold for somatic spikes | 18mV |
| $\tau_{adapt}$ | Adaptation time constant of excitatory neurons | Baseline excitability: 200msec<br>High excitability: 100msec |
| $\beta_{adapt}$ | Adaptation reset constant | Baseline excitability: 9<br>High excitability: 4 |
| $A_{adapt}$ | Adaptation coupling parameter | 0.02 |
| $\tau_{bAP}$ | Back propagating action potential time constant | 10msec |
| $E_{bAP}$ | Back propagating action potential max amplitude | 5 mV |
| $\Theta_{PRP}$ | Calcium threshold for somatic Plasticity-Related Protein (PRP) synthesis | 40.0 |
| $\tau_{PRP}$ | Time constant for PRP decay | 60 minutes |
| $\tau_H$ | Time constant of homeostatic synaptic scaling | 720 hours |

**Suppl. Table 2: Parameters of the microcircuit model**
